# The molecular mechanism of cytoadherence to placenta or tumor cells through VAR2CSA from *Plasmodium falciparum*

**DOI:** 10.1101/2021.07.06.451098

**Authors:** Weiwei Wang, Zhaoning Wang, Xiuna Yang, Yan Gao, Xiang Zhang, Weiwei Wang, Lingyu Zhao, Jun zhang, Qingqing Jing, Qian Xu, Long Cao, Yuan Tian, Aguang Dai, Jin Sun, Lei Sun, Lubin Jiang, Zhenguo Chen, Lanfeng Wang

**Affiliations:** The Center for Microbes, Development and Health, CAS Key Laboratory of Molecular Virology & Immunology, Institut Pasteur of Shanghai, Chinese Academy of Sciences; University of Chinese Academy of Sciences, Shanghai, China; College of Life Sciences, University of Chinese Academy of Sciences, 100049 Beijing, China; Shanghai Fifth People’s Hospital, Fudan University, and Shanghai Key Laboratory of Medical Epigenetics, International Co-laboratory of Medical Epigenetics and Metabolism (Ministry of Science and Technology), Institutes of Biomedical Sciences, Fudan University, Shanghai, China; Institute for Advanced Immunochemical Studies and School of Life Science and Technology, ShanghaiTech University, Shanghai, China

**Keywords:** VAR2CSA, CSA, Cryo-EM structure, PAM, Cytoadherence

## Abstract

Pregnancy Associated Malaria (PAM) threatens more than one million pregnant women and their infants in endemic regions due to poor outcomes. VAR2CSA plays a vital role in the cytoadherence of infected erythrocytes (IEs) to placenta against immuno-clearance via binding to Chondroitin sulfate A (CSA), which is displayed mostly on the surface of placental or tumor cells. In this study, we determined the cryo-EM structures of VAR2CSA ectodomain and its complex with CSA at a resolution of 3.6 Å and 3.4 Å, respectively and revealed that CSA binding induces significant conformational change for ligand accommodation. Beyond structural studies, we generated VAR2CSA fragments and mutation and evaluated their binding activity to either isolated CSA or placental and tumor cell line using multi-disciplinary techniques. The results showed that 9-site mutation in DBL2X abolished the CSA binding activity and also disrupted its binding to both placental and tumor cells. Overall, our work clearly elucidated the molecular mechanism of cytoadherence to placental or tumor cells through VAR2CSA. Which may facilitate new PAM vaccine development and improve the drug delivery systems targeting both placenta and tumors.

## Introduction

Malaria, a mosquito-borne infectious disease, is caused by a single-cell protozoan named as *Plasmodium*, of which *P.falciparum* is the most virulent species^1^. Since women in pregnancy are more vulnerable to *Plasmodium* infection, PAM attracts more attention due to the poor outcomes including maternal anemia, stillbirth, preterm delivery, infant death, *etc*^2,3^. The World Health Organization (WHO) reported that there are more than one million women suffering from PAM in endemic regions in 2019^4^.

In the pathogenesis of PAM, IEs could be sequestated in the intervillous space of the placenta to avoid immune clearance. VAR2CSA, a *P. falciparum* encoded protein expressed on the suface of IEs, plays a vital role in cytoadherence and sequestration ^5,6^. VAR2CSA belonging to *P. falciparum* erythrocyte membrane protein 1 (PfEMP1) family is a 350 kDa transmembrane protein comprising of N-terminal etcodomain (~306 kDa), transmembrane domain, and C-terminal cytoplasmic domain^7,8^. In terms of composition, the ectodomain can be divided into six Duffy-binding-like (DBL) domains (DBL1X, DBL2X, DBL3X, DBL4**ε**, DBL5**ε**, DBL6**ε**), N-terminal sequence (NTS), and multiple inter-domains (IDs) (ID1, ID2a, ID2b, ID3)^9–11^. It was proposed that the ectodomain is responsible for specifically binding to the CSA displayed mainly in placenta. Chondroitin sulfate (CS), a sulfated glycosaminoglycan, is mostly found attached to proteins as a type of glycosylation with complex saccharides sulfated in various positions and quantities. Additionally, CSA is a specific glycosaminoglycan with the C4 of the N-acetyl-D-galactosamine (GalNAc) sulfated. Moreover, a dodecasaccharide with six disaccharide repeats (GalNAc and glucuronic acid (GlcA)) in low sulfated form has been considered as the minimal structure motif in CSA for IEs’ adherence^10,11^. It has been reported that CSA presented in the syncytiotrophoblast layer of pregnant women is considered different from those found in other normal tissues. In addition, CSA has been widely identified in solid tumors mostly due to the similar phenotype between tumors and placenta^12^.

VAR2CSA initiates the pathogenesis of PAM through recognizing and binding CSA, thus is widely considered as a leading potential vaccine antigen to elicit protective immune response or a perspective drug target against PAM. The preparation of VAR2CSA full length or ectodomain with high stability and yield is a sustained challenge due to the large size and complex architecture in spite of a couple of successful reports^13–15^. This issue significantly hampers understanding its structure and function, then developing effective vaccine candidates as well. Surprisingly, almost all individual DBL domains with different borders have been shown to induce inhibitory antibodies to the cytoadherence of IEs^16–21^. Moreover, naturally acquired antibodies showing high inhibitory activity other than those elicited by individual recombinant VAR2CSA fragment suggests that ideal antigens might be the combination of different VAR2CSA fragments^22^. Currently, two vaccine candidates derived from VAR2CSA (PRIMVAC and PAMVAC) showed reasonable safety and immunogenicity in Phase 1 clinical trials^23,24^. Additionally, CSA binding peptide derived from VAR2CSA can be applied to placenta targeted drug delivery system^25^. Due to the wide distribution of CSA in various tumors, either recombinant fragment or 28-residue polypeptide from VAR2CSA has been showed to have the potential for effective delivery of specific chemicals targeting various tumors^12,26^. Meanwhile, recombinant VAR2CSA shows promising clinic application in pre-diagnosis of malignant tumors through specific binding to circulated tumor cells^27,28^.

So far, a few structures of VAR2CSA fragments (i.e., DBL3X, DBL3X-DBL4ε, DBL6ε) have been solved^29–34^. Additionally, the structure model from the reconstruction of VAR2CSA ectodomain using negative staining electron microscopy proposed an overall domain arrangement^35^. We are still lack of sufficient structure information to understand the architecture and ligand binding mechanism of VAR2CSA.

In this study, using cryo-electron microscopy single-particle analysis, we determined the three-dimensional structures of *P. falciparum* VAR2CSA ectodomain and its complex with CSA at a resolution of 3.6Å and 3.4 Å, respectively. Structural analysis identified that a dodecasaccharide with six sulfated disaccharide repeats from CSA is located at the binding pocket formed by N-terminal sequence (NTS), DBL1X, DBL2X, and DBL4ε. Intriguingly, it was revealed that CSA binding induces obvious conformational change to close the binding pocket by turning DBL2X and DBL1X closer to DBL4ε, and meanwhile enlarge the inner binding pocket via slightly moving a CSA-binding helix of DBL2X outward. In addition, the structural analysis indicated that 9 key residues with positive charge in DBL2X might be mainly responsible for CSA binding, which is further validated by sequence alignment, site mutagenesis, and CSA binding tests. Most importantly, we tested the binding activity of various VAR2CSA fragments and mutants to both placental cells and tumor cells using confocal microscopy. The result regarding the CSA binding mechanism is fully in line with our biochemistry results above. In summary, we elucidated the detailed molecular mechanism of VAR2CSA recognizing and binding CSA displayed on the surface of both placental cells and tumor cells using multi-disciplinary techniques.

## Results

### Cryo-EM structure of VAR2CSA ectodomain

Due to the large complexity of VAR2CSA, the preparation of full length or ectodomain protein with high quality is a major issue for understanding the structure and function. In this study, VAR2CSA ectodomain (residue1-2630, Fig. 1A) coding DNA fragment was inserted into a secretory vector gp67-438-B modified based on 438-B vector. Sf9 insect cells infected by recombinant baculovirus harboring interest gene were cultured in suspension serum free system to express VAR2CSA ectodomain, which was purified through ammonium sulfate precipitation and sequential column chromatography.

**Fig. 1.**
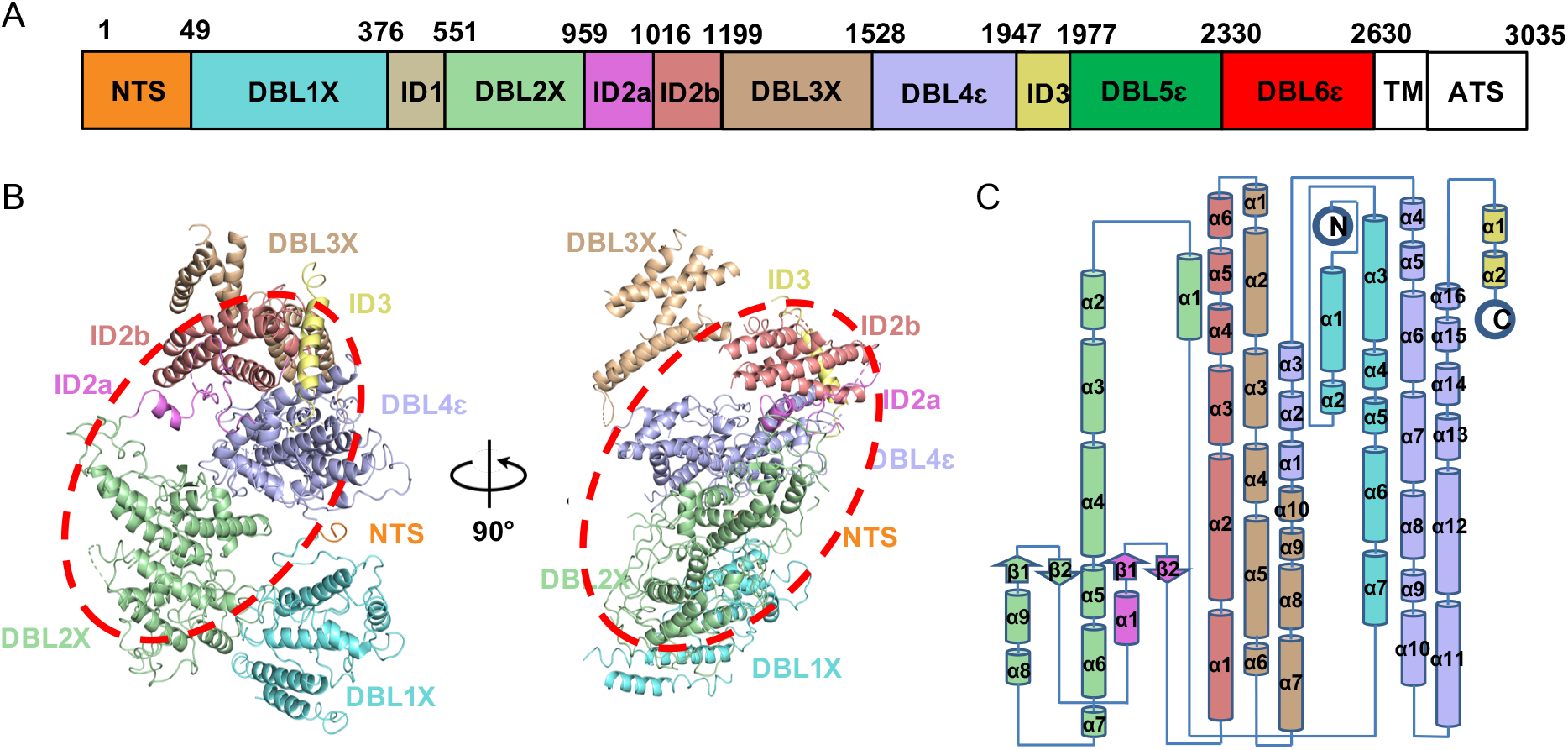
Overall structure of apo VAR2CSA ectodomain. **A**, Schematic sequence architecture of VAR2CSA of *Plasmodium falciparum* 3D7 strain. Domains reconstructed according to the cryo-EM density map are colored respectively. NTS, orange; DBL1X, aquamarine; DBL2X, pale green; ID2a, violet; ID2b, salmon; DBL3X, wheat; DBL4ε, light blue; ID3, pale yellow; DBL5ε, green; DBL6ε, red; ID1; grey. Color codes are used throughout this paper unless otherwise noted. **B**, The ribbon diagram of apo VAR2CSA ectodomain core region shown in two different views; The core center highlighted using red dashed oval. **C**, The topology of apo VAR2CSA ectodomain core region. α-Helixes shown as cylinders; β-strands shown as arrows.

The analysis using size exclusion chromatography along with reduced and non-reduced SDS-PAGE showed that VAR2CSA ectodomain was mostly monomer, however there was still a small fraction of dimer stabilized by intermolecular disulfate bond (Fig. S1A). Since there are 114 cysteine residues in the VAR2CSA ectodomain, the inter or intra molecular disulfate bonds may affect the conformation and stability. So, we tried to stabilize VAR2CSA ectodomain at certain conformation using glutaraldehyde Gradient Fixation (Grafix). The presence of both monomer and dimer in above crosslinked VAR2CSA ectodomain was further confirmed by SDS-PAGE (Fig. S1B). Both monomer and dimer were imaged using negative staining and transmission electron microscopy (TEM). It was shown that the monomer seems like a revolver in 2D classification with a solid center and flexible ends(Fig. S2A), which was mostly in agreement with the previous report^35^. Intriguingly, the dimer looks like a butterfly in 2D classification and low-resolution model (Fig. S2B).

Next, the monomer was applied to single particle analysis using cryo-EM. The structure of VAR2CSA ectodomain was determined at an overall resolution of 3.6 Å (Fig. 1B, Fig. S3/S4). Intriguingly, a part of clear density connected to core region could be visualized in the cryo-EM map under low threshold and thus named as wing region (Fig. S3C/S4A). Due to the high flexibility of wing region (Fig. S3C), we cannot get high resolution map to build atomic model, so we focused on the core region for model building and structural analysis. We started model building using Phenix.map_to_model to generate an initial model with helixes fitting well in density^36^. Then we tried *de novo* model building using the published structure of DBL4ε (PDB ID: 4P1T) as a reference^32^, because the center of core region has higher resolution up to 3.1 Å, where the side chains are clearly tracible. As resolution decreases, structural model was built from the center to the periphery. In line with the proposed domain arrangement^35^, our structure showed that the core region covers NTS, DBL1X, DBL2X, ID2a, ID2b, DBL3X, ID3, and DBL4ε (Fig. 1B/1C). Meanwhile, DBL5ε and DBL6ε form wing region, which is too flexible to be clearly interpreted currently. As for the core region, DBL2X and DBL4ε stacking closely with ID2a, ID2b, and ID3 form the most stable core center, which serves as a base for anchoring DBL3X and DBL1X at top or bottom site, respectively (Fig. 1B).

### Cryo-EM structure of VAR2CSA ectodomain complex with CSA

We next want to know how VAR2CSA ectodomain recognizes and binds to CSA from the structural perspective. Firstly, three potential VAR2CSA interacting glycoproteins (Podocalyxin-like protein 2 (PODXL2), Cluster of Differentiation 44 (CD44), and Decorin)^12^ recombinantly expressed in HEK 293 cells and CSA^37^ from bovine trachea (bought from Sigma) were prepared to interact with VAR2CSA ectodomain. We incubated VAR2CSA ectodomain with the substrates above and evaluated the interaction using gel filtration column, individually. The results showed that only CSA can form a stable complex with VAR2CSA ectodomain (Fig. 2A). However, the affinity of those three glycoproteins expressed *in vitro* is generally low, so there are no obvious complex peaks detected in chromatography (Fig. S5).

**Fig. 2.**
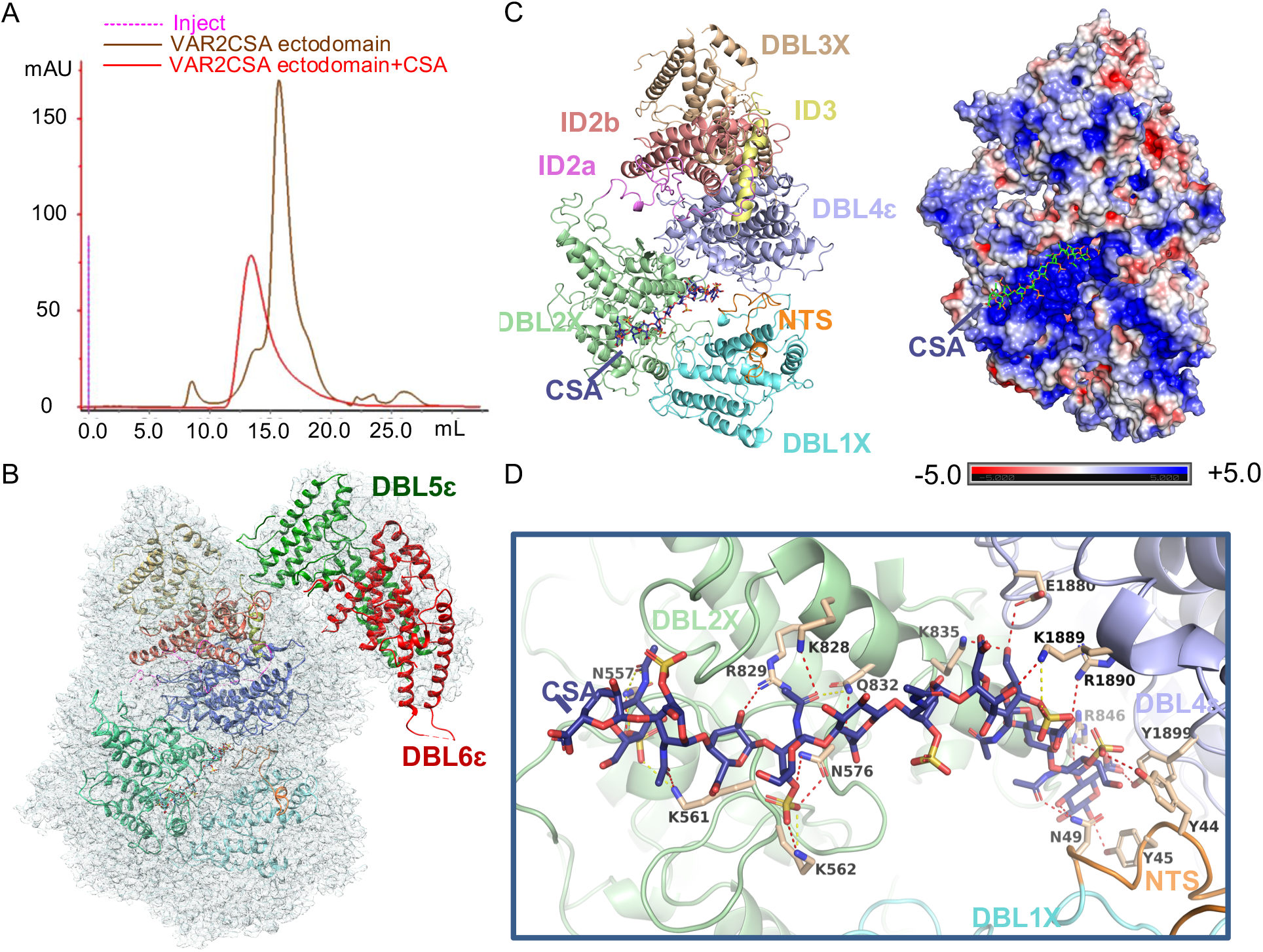
Overall structure of VAR2CSA ectodomain complex with CSA. **A**, The comparison of gel filtration chromatography between apo VAR2CSA ectodomain and its complex with CSA (VAR2CSA-CSA) on a Superose 6 increase 10/300 GL column. **B**, Density map of VAR2CSA-CSA with final structure model represented by ribbon diagram. **C**, The ribbon diagram (Left) and surface electrostatic potential map (Right) of VAR2CSA-CSA core region. CSA drawn as sticks. **D**, Interactions between CSA and VAR2CSA were shown in the binding pocket. Interacting residues from NTS, DBL2X, and DBL4ε were highlighted individually.

The purified VAR2CSA ectodomain complex with CSA (VAR2CSA-CSA) was used to prepare cryo-sample for screening and data collection via both 200kV and 300kV TEM. The structure of VAR2CSA-CSA was solved at an overall resolution of 3.4 Å (Fig. 2B/C, Fig. S6/S7). Compared with the apo structure, densities of the wing region have been significantly improved, which makes it feasible for the flexible fitting of both DBL5ε and DBL6ε. (Fig. 2B). As for the core region, the overall domain architecture is similar to that of apo structure. Briefly, DBL2X and DBL4ε along with ID2a, ID2b, and ID3 form the stable core center serving as a base for anchoring DBL3X and DBL1X at either site. Most importantly, a highly basic pocket formed by DBL2X, DBL1X, NTS, and DBL4ε has been identified to accommodate a dodecasaccharide with six sulfated disaccharide repeats from CSA, which could be well fitted into the density map (Fig. 2C, Fig. S8B). Intriguingly, 16 residues were identified to be responsible for the direct interaction with 10 of 12 monosaccharides except the 6^th^ and 11^th^ unit of CSA dodecasaccharide. Among them, there are 9, 4, or 3 residues derived from DBL2X, DBL4ε, or NTS, respectively. Moreover, the 9 residues of DBL2X were shown to interact directly with 7 monosaccharide units (Fig. 2D, Table S1).

### Significant conformational change observed upon CSA binding

When we compared the density maps of VAR2CSA ectodomain in the presence or absence of CSA using either 200 kV or 300 kV cryo-EM datasets, it was shown that there is a significant conformational change induced by CSA binding in the VAR2CSA core region, especially for the DBL1X domain (Fig. S9). Moreover, when we tried to build the model of VAR2CSA-CSA using our apo structure as a start reference, rigid body refinement did not work quite well. That also indicated that the conformational change might be induced upon CSA binding. Due to the high resolution and potential high stability in the core center, DBL4ε was used as immobilized reference for the structure alignment between VAR2CSA-CSA and VAR2CSA ectodomain (Fig. 3). Interestingly, conformational change in core areas has been further confirmed as following (Video 1). Firstly, there is 1.8 Å outward bent for the key CSA binding helix of DBL2X in deep pocket to make enough space to accommodate the ligand (Fig. 3A). Meanwhile, the rest part of DBL2X move about 2 Å toward the pocket (Fig. 3B). Most significantly, the DBL1X moves closer to DBL4ε at more than 3.2 Å (range from 3.2 Å to 4.4 Å) for multi-helixes to facilitate NTS interacting with CSA and close the pocket as well (Fig. 3C). Additionally, DBL3X also has a big conformational change outward mostly due to high flexibility (Fig. 3D).

**Fig. 3.**
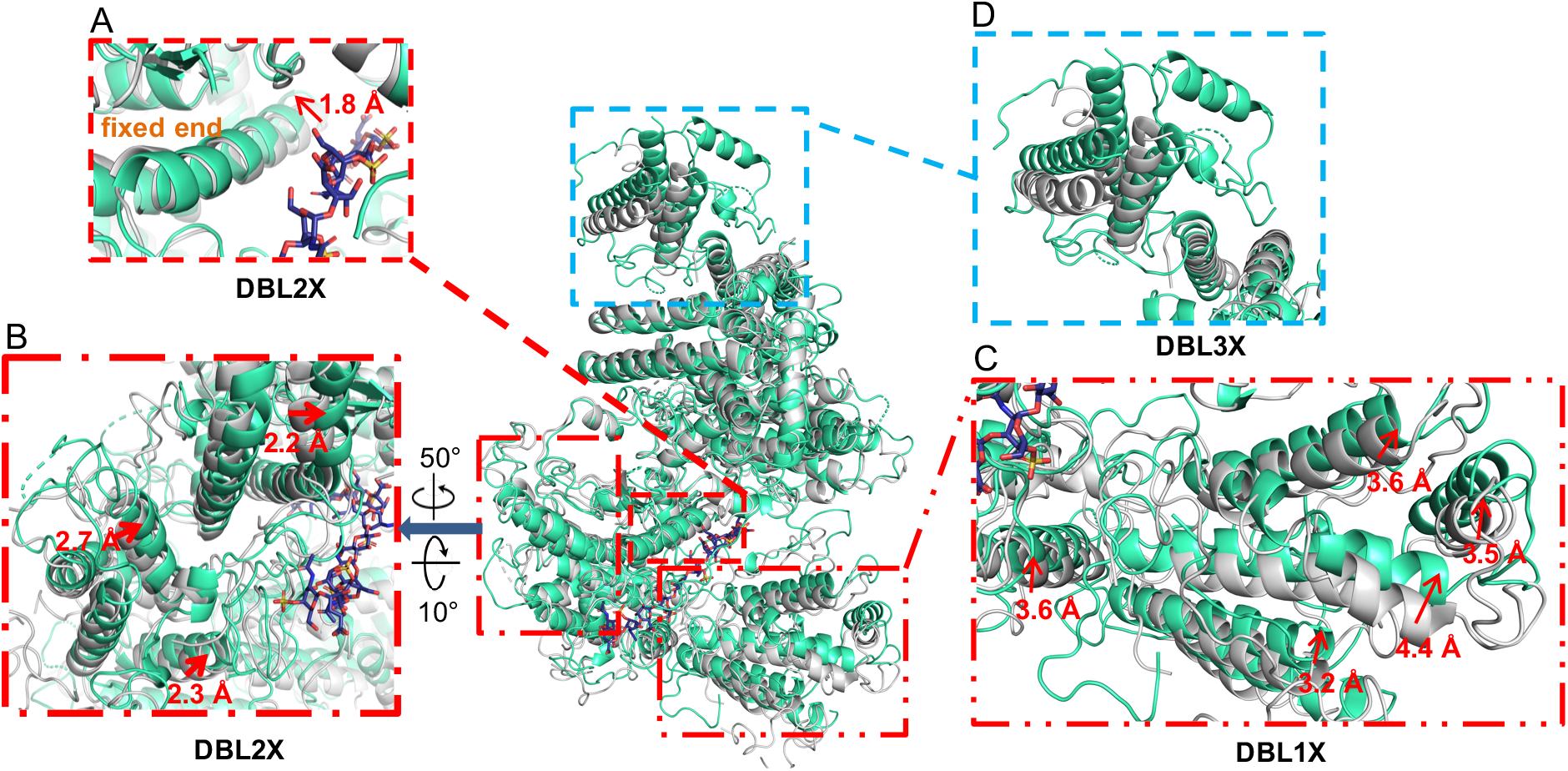
Structural Comparison between VAR2CSA-CSA with apo VAR2CSA ectodomain by aligning DBL4ε domains. **A**, The helix associated with CSA in DBL2X domain exhibited an 1.8 Å bent outward at the binding site while the distal end remained fixed. **B**, The surficial helixes in DBL2X move about 2 Å toward the CSA binding pocket. **C**, The DBL1X domain went through a significant conformational change at a distance of more than 3.2 Å to close the CSA binding pocket. **D**, Conformational change was shown in DBL3X domain.

### DBL2X plays an essential role in binding CSA

Although the CSA binding pocket has been clearly revealed in our complex structure, the identification of the minimal structural elements for CSA binding is of importance for rational design of PAM vaccine antigens and tumor targeted application as well. The structural analysis showed that compared with NTS or DBL4ε, DBL2X has the most residues (9 aa) directly interacting with 7 out of 10 VAR2CSA-associated monosaccharides. Subsequently, we evaluated whether DBL2X plays an essential role for binding CSA. At a first glance, the individual structures of four DBL domains (DBL1X, DBL2X, DBL3X, and DBL4ε) are quite similar to each other, with the typical “3+2” helix bundle mode shared (Fig. 4A). However, the sequence alignment of 6 individual DBL domains showed that the 9 key residues in DBL2X are rarely conserved, which indicates that DBL2X might be mainly responsible for CSA binding other than the rest DBL domains (Fig. 4B, Fig. S10). In addition, when we aligned the DBL2X sequences of VAR2CSA across various *P. falciparum* strains, those 9 residues are highly conserved, which also suggests a common CSA recognition mechanism dominated by DBL2X (Fig. 4B, Fig. S11). Next, we generated two VAR2CSA truncations: VAR2CSA-50-962 (DBL1X and DBL2X) and VAR2CSA-550-962 (DBL2X alone). Then the CSA binding activity of both truncations was evaluated using gel filtration chromatography. The results showed that similar with VAR2CSA ectodomain, both truncated proteins could form stable complex with CSA (Fig. 5A), which is in line with the previous report that DBL2X might be the minimal requirement for binding CSA^38^.

**Fig. 4.**
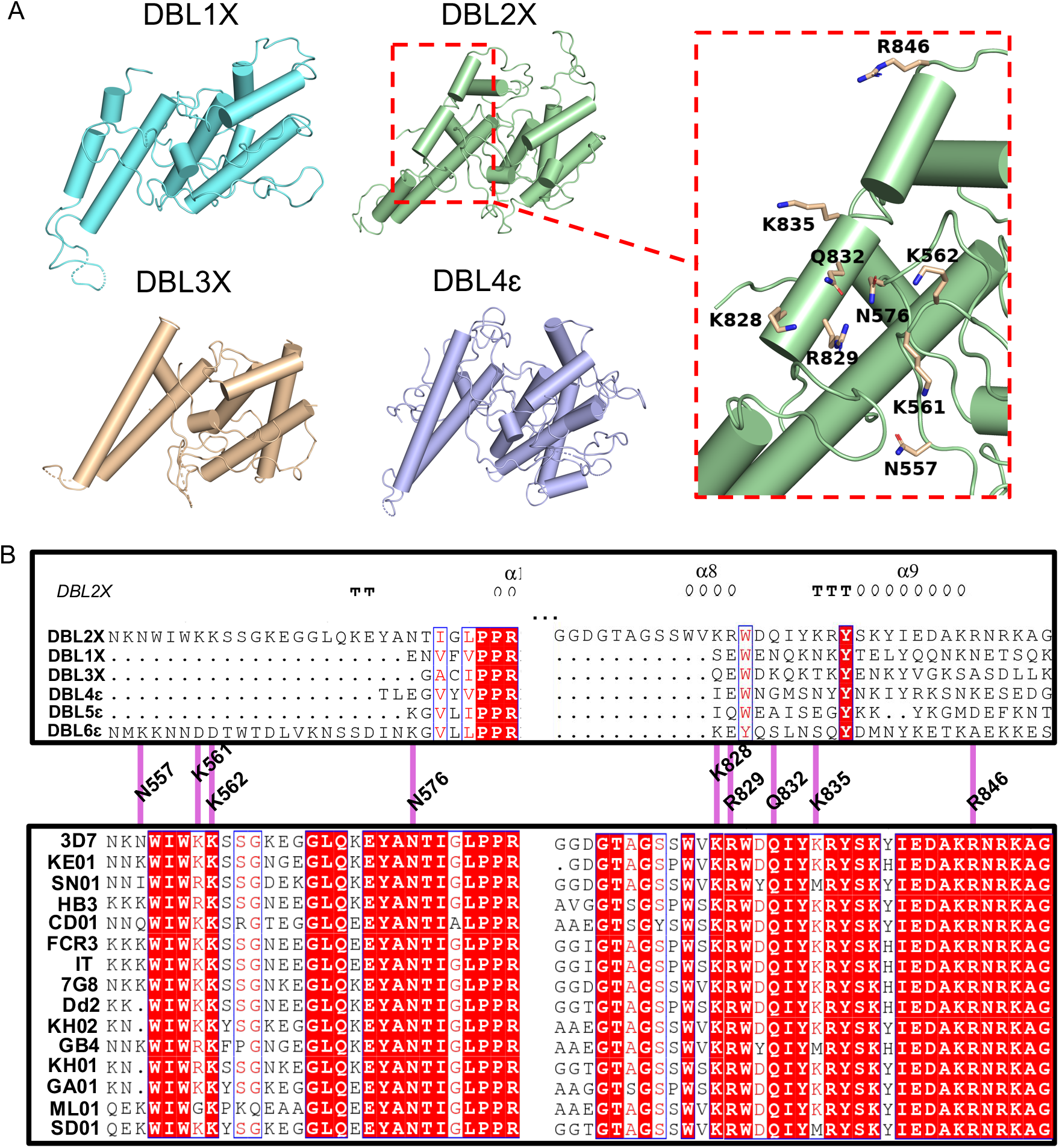
The key residues in DBL2 are responsible for CSA binding *in vitro*. **A**, The overall structures of four individual DBL domains of the core region were shown. The nine CSA binding residues in DBL2X were highlighted. **B**, Sequence alignment of the DBL2X domain with the rest VAR2CSA DBL domains (Up) or with those DBL2X domains from different *P. falciparum* species (Down) as well. The nine CSA binding residues in DBL2X were indicated by magenta lines.

**Fig. 5.**
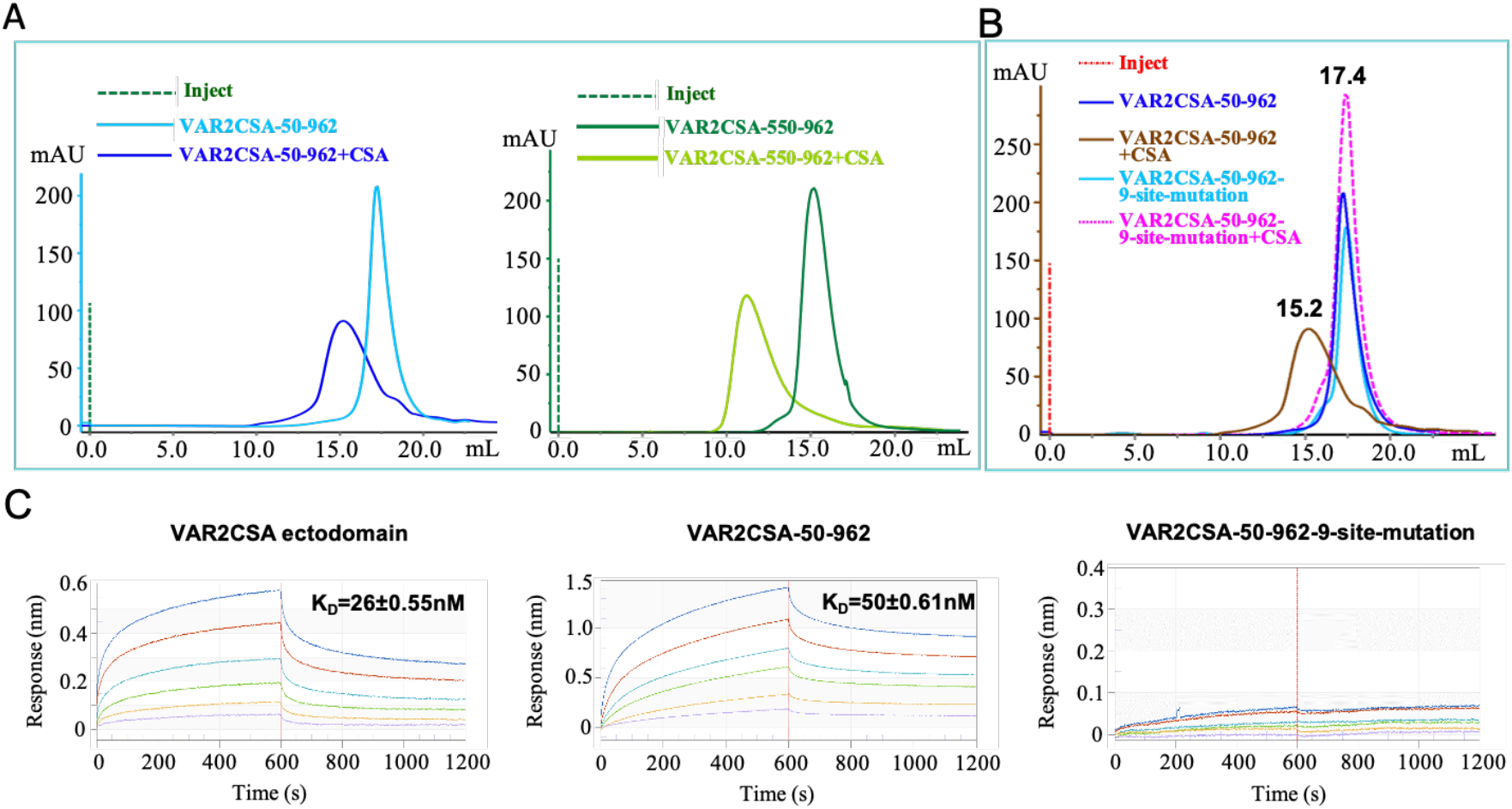
The mutation of the 9 key residues in DBL2X can abolish the CSA binding activity of VAR2CSA *in vitro*. **A**, Gel filtration chromatography comparison of VAR2CSA-50-962 (including DBL1X and DBL2X, left) and VAR2CSA-550-962 (DBL2X, right) in the presence or absence of CSA. Left, Superose 6 Increase 10/300 GL column; Right, Superdex75 10/300 GL column. **B**, Gel filtration chromatography of VAR2CSA-50-962 and VAR2CSA-50-962-9-site-mutation (N557D, K561E, K562E, N576D, K828E, R829E, Q832E, K835E, R846E) in the presence or absence of CSA on a Superose 6 increase 10/300 GL column. **C**, The binding kinetics of VAR2CSA ectodomain, VAR2CSA-50-962, and VAR2CSA-50-962-9-site-mutation were further assessed by BLI assays, and equilibrium constant (K_D_) values were calculated accordingly.

### 9-site mutagenesis in DBL2X abolished the CSA binding capacity

Since VAR2CSA ectodomain, VAR2CSA-50-962, and VAR2CSA550-962 all could form stable complex with CSA in gel filtration chromatography (Fig. 2A, Fig. 5A), VAR2CSA-50-962 was selected as a representative for the following experiments. The mutation with 9 key residues mutated to D or E (N557D, K561E, K562E, N576D, K828E, R829E, Q832E, K835E, R846E) was named as VAR2CSA-50-962-9-mutation. Then both the wild type and mutant of VAR2CSA-50-962 in the presence or absence of CSA were subject to the analysis using gel filtration chromatography on a Superose 6 increase 10/300 GL column. The chromatogram comparison showed that in the absence of CSA, the retention volume of VAR2CSA-50-962-9-mutation is almost the same as that of the wild type. More importantly, in the presence of CSA, the retention volume of the mutant remained unchanged, however the retention volume of wild type VAR2CSA-50-962 was significantly shifted due to the formation of stable super-complex (Fig. 5B). Additionally, VAR2CSA ectodomain, VAR2CSA-50-962, and VAR2CSA-50-962-9-mutation were used to test their affinity to CSA by Octet RED 96. In accordance with the results of chromatography above, VAR2CSA ectodomain and VAR2CSA-50-962 exhibited strong affinity to CSA, while VAR2CSA-50-962-9-mutation showed totally no binding activity to CSA under test conditions (Fig. 5C). The results above indicated that the 9-site mutation abolished the CSA binding activity of VAR2CSA-50-962 *in vitro*.

### 9-site mutagenesis in DBL2X disrupted cytoadherence to placental and tumor cells

Next, whether the 9-site mutation could influence the binding activity of VAR2CSA to placental cells or tumor cells was further verified using confocal fluorescence microscopy, respectively. We firstly tested the binding activity of VAR2CSA ectodomain, VAR2CSA-50-962, and VAR2CSA-50-962-9-site-mutation to JEG-3 cells as a representative of placental cells. Briefly, 500 nM individual protein was incubated with JEG-3 cells preseeded on a slide sufficiently. The slides were then washed by three times and unbound proteins were thus removed thoroughly. The fluorescence labeled anti-His monoclonal antibody was supplemented to stain the recombinant proteins above, while DAPI was applied to stain the cell nuclei. Then, the slides were subjected to analysis using Olympus FV1200 Laser scanning confocal microscope to test the binding activities of VAR2CSA fragments and mutant to JEG-3 cells. Interestingly, it was shown that compared with mock (No proteins added), VAR2CSA-50-962 and VAR2CSA ectodomain all could specifically bind to JEG-3 cells (Fig. 6A). Moreover, no obvious difference of the binding affinity was observed among these two proteins. Strikingly, the cell binding activity of VAR2CSA-50-962-9-site-mutation was mostly abolished (Fig. 6A). Meanwhile, HepG2 as a representative of tumor cells has also been used to test the cytoadherence of VAR2CSA fragments above via confocal fluorescence microscopy. Roughly, HepG2 cell samples for confocal fluorescence microscopy were prepared following the same procedures as described above for JEG-3 cells. The results showed that both VAR2CSA-50-962 and VAR2CSA ectodomain have strong cytoadherence to HepG2 cells (Fig. 6B). However, VAR2CSA-50-962-9-site-mutation is almost lack of the ability of cytoadherence to HepG2 cells (Fig. 6B). That is fully in line with the biochemistry results above.

**Fig. 6.**
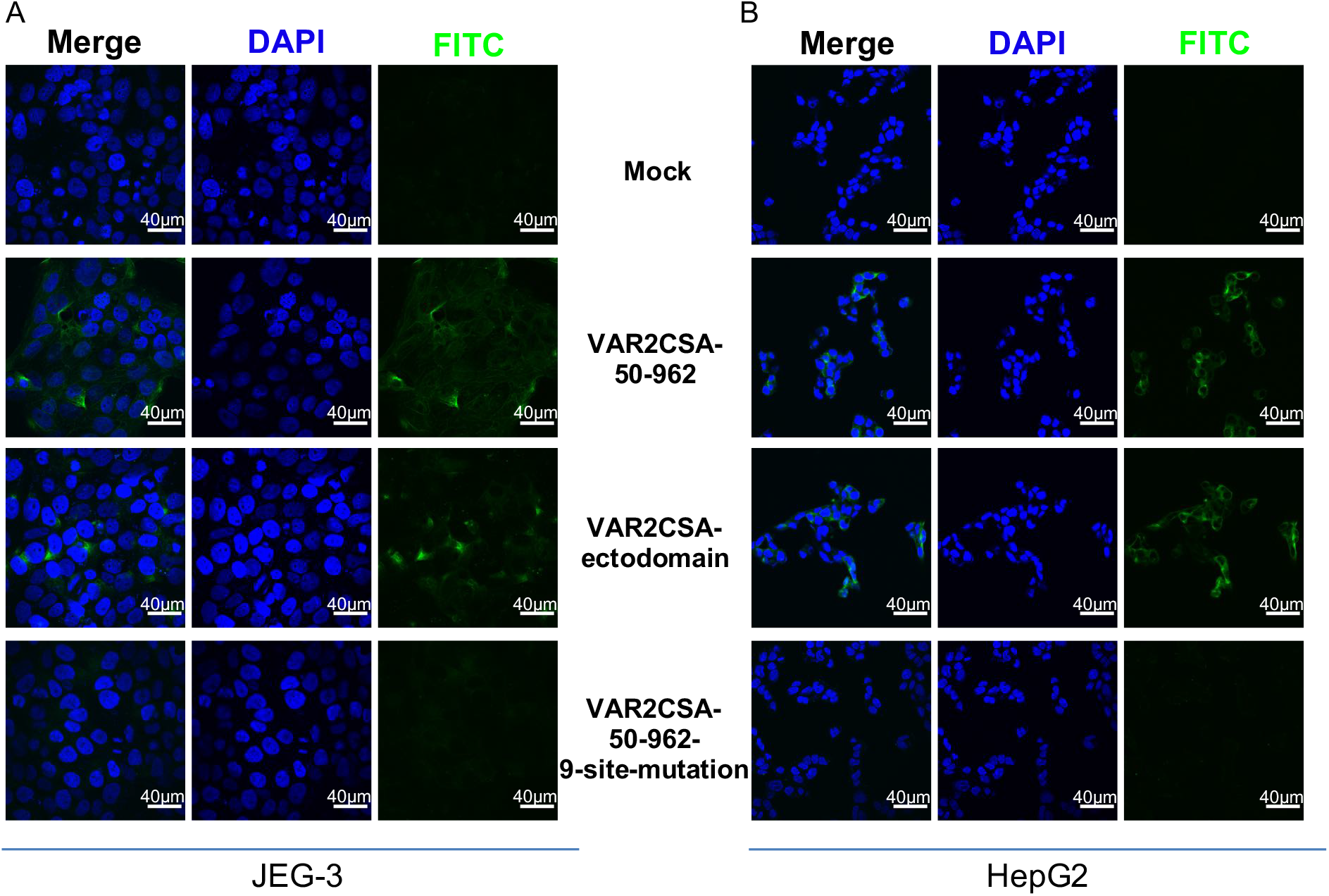
The mutation of the 9 key residues in DBL2X can disrupt VAR2CSA binding to placental or tumor cells. The binding activities of VAR2CSA-50-962, VAR2CSA ectodomain, and VAR2CSA-50-962-9-site-mutation to JEG-3 cells (**A**) or HepG2 cells (**B**) were tested by immunofluorescence assay using confocal fluorescence microscopy, respectively. Green (stained with Anti-His-FITC antibody) indicates VAR2CSA proteins; blue (DAPI) represents cell nuclei.

Taken together, the 9-site mutation in DBL2 can abolish the binding activity of VAR2CSA to either placental cells or tumor cells under test conditions.

### Work model clarifying the mechanism of cytoadherence to placenta or tumor cells through VAR2CSA

Based on the above information acquired from structure, site mutagenesis, gel filtration chromatography, and confocal fluorescence microscopy, we made a cartoon model to simply clarify the functional mechanism of VAR2CSA. In terms of CSA binding, VAR2CSA designed as a cannibal plant recognizes and binds to CSA drawn as tiny zombie through a giant mouth formed by DBL2X, DBL1X, NTS, and DBL4ε. Upon zombie entering, the big jaw standing for DBL1X will close to tightly catch the quarry (Fig. 7A). In the cell environment, distinguished from specific CSA modification, other glycosylations were drawn as sunflowers. Meanwhile tumor cells, placental cells, and other cells were shown as balloons in salmon, pale yellow, and pale green, respectively. According to the same mechanism proposed above, the cartoon further shows VAR2CSA characteristically recognizing and binding either tumor cells or placental cells instead of other normal cells (Fig. 7B).

**Fig. 7.**
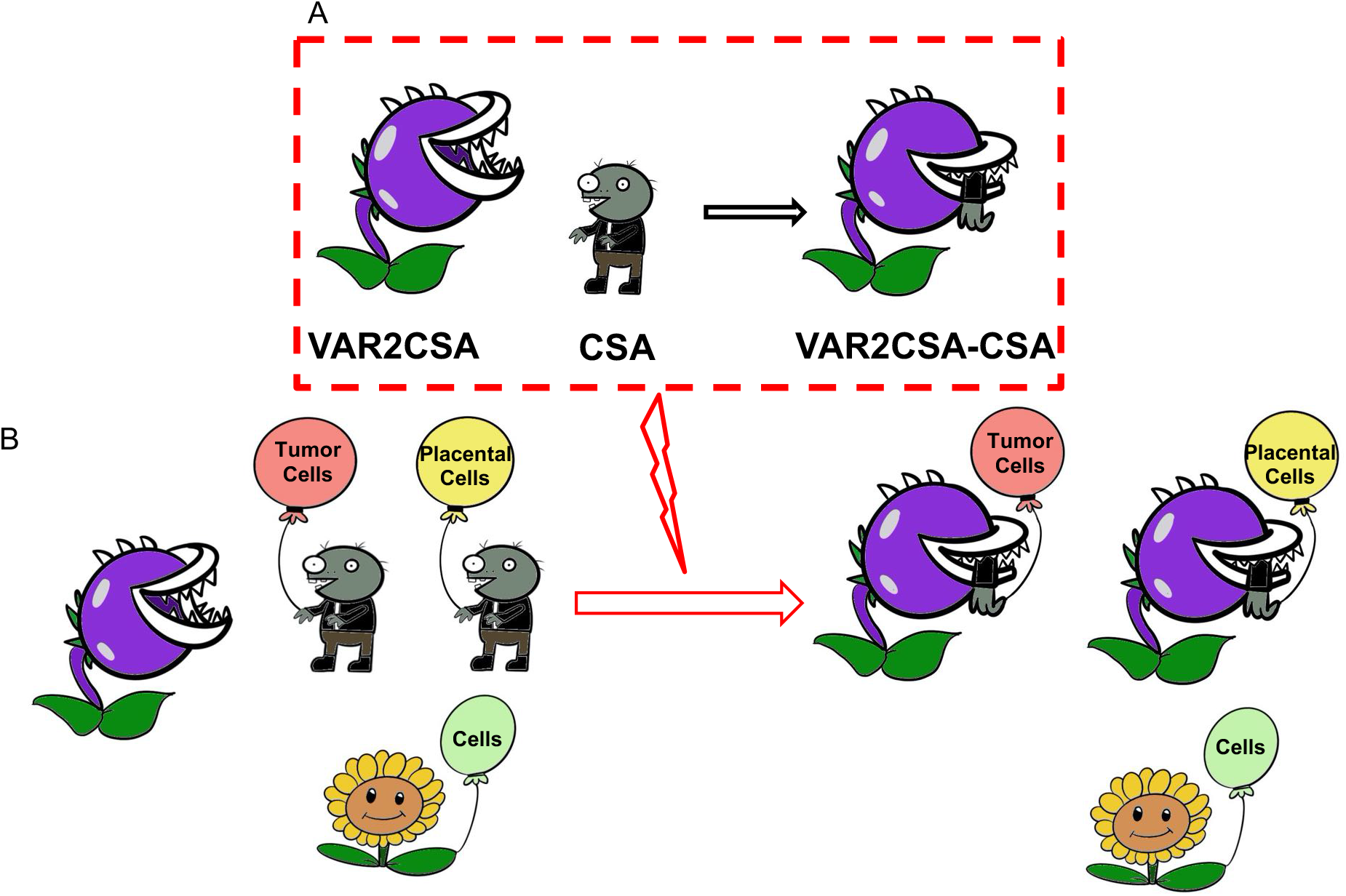
Model of VAR2CSA recognizing both Placental cells and Tumor cells. **A**, The cartoon simply exhibits the mechanism of VAR2CSA capturing CSA ligand and reveals the conformational change upon CSA binding. **B**, The cartoon further shows the mechanisms of VAR2CSA characteristically recognizing either placental cells or tumor cells instead of other normal cells by recognizing the specific CSA displayed on the cell surface. VAR2CSA were designed as cannibal plants, with two states corresponding to CSA binding or not, respectively. CSA modification were drawn as cute zombies; other glycol-modifications as sunflowers. Cells were shown as balloons, indicated in different colors. Tumor cells, salmon; Placental cells, yellow; Other normal cells, light green.

## Discussion

VAR2CSA acts as one of the most important factors in the pathogenesis of PAM^5,6^. Determination of the architecture of VAR2CSA is essential to understand how it functions. Sequence analysis showed that 6 DBL domains sharing high conservation constitute the skeleton of VAR2CSA. Although a few crystal structures of DBL domains have been solved^29–34^, there is little knowledge about the detailed architecture and the functional mechanism of VAR2CSA, except one proposed model^35^. In our opinion, the most challenging part is that it is hard to obtain functional VAR2CSA with high yield and quality due to the large size and complexity. Even though, a couple of groups reported that VAR2CSA ectodomain wild type or mutant with potential N-glycosylation sites substituted with alanine have been expressed in either mammalian or insect cells^13–15^. In this study, we modified the expression and purification strategy to gain VAR2CSA ectodomain with high yield (up to 3 mg/L) and quality, which significantly speeds up the structural and functional study. In the long run, it also facilitates the large-scale preparation of VAR2CSA ectodomain as effective antigen candidates with higher immunogenicity compared with shorter fragmented antigens previously reported^14,15^.

Most importantly, significant conformational change in VAR2CSA upon CSA binding has been firstly described in this study. We compared the structures of both VAR2CSA ectodomain and VAR2CSA-CSA through overlaying the most stable DBL4ε domain. The result showed that CSA binding pushes the interacting motif of DBL2X outward in the deeper binding pocket by 1.8 Å to get more space for accommodation. Meanwhile, the surficial part of DBL2X moves inward around 2.0 Å to slightly shrink the pocket. Most strikingly, DBL1X moves closer to DBL4ε by more than 3.2 Å to close the binding pocket and assist NTS to interact with CSA. The conformational change above is mostly in line with the so-called “induced-fit model”^39^, commonly accepted for certain enzymes binding ligands. It may also help to interpret the sequestration of infected IEs through this special tight binding mechanism.

Intriguingly, all key interacting residues within 3.5 Å relative to CSA were divided into 3 groups according to their domain affiliation, among which DBL2X with 9 key residues accounts for the most contribution to CSA binding. The 9 key residues in DBL2X are less conserved between 6 DBL domains but highly conserved across different *P.falciparum* strains. In agreement with previous reports^38,40,41^, VAR2CSA truncations (50-962 and 550-962) containing DBL2X were shown to form stable complexes with CSA, similar as VAR2CSA ectodomain. However, 9-site mutation in VAR2CSA-50-962 abolished the formation of stable complex with CSA compared with wild type. Most improtantly, the confocal fluorescence microscopy further showed that VAR2CSA ectodomain, VAR2CSA-50-962 can specifically bind not only to JEG-3 cells as a representative of placental cells but also to HepG2 cells as a representative of tumor cells with comparable affinity, however, 9-site mutation in VAR2CSA-50-962 abolished its binding activity to both cells. Altogether, our biochemistry and cell biology experiments showed for the first time that 9 key residues in DBL2X mainly account for the CSA binding and the seqestration of both placental cells and tumor cells.

Specific glycosylation in proteoglycans is the decisive factor recognized by VAR2CSA. As reported previously, CSA from placental or tumor cells is optimal for binding VAR2CSA^12^. We measured the interaction between VAR2CSA ectodomain and a few potentially interacting glycoproteins (i.e., PODXL2, CD44, and Decorin) recombinantly expressed in HEK293 expression system or CSA commercially available from Sigma^37^, respectively. The results showed that only CSA can interact with VAR2CSA ectodomain to form a stable complex, however, the affinity of the glycoproteins expressed *in vitro* is generally low. That indicated the glycosylated modification type of proteoglycans plays a decisive role in its ability to interact with VAR2CSA. Meanwhile it suggests that tissuespecific glycosylation-related enzymes might be responsible for CSA synthesis in either placental cells or tumor cells.

Interestingly, there are more than 100 cysteines capable of forming inter or intra molecular disulfate bonds in the VAR2CSA ectodomain, which could stabilize individual proteins, or maintain the complexity *in vivo* under various reduced or oxidized environments. Our result showed that the recombinant VAR2CSA ectodomain behaves mainly as a monomer, however a portion of homodimer has also been detected and further confirmed using Grafix. Most intriguingly, the homodimer of VAR2CSA ectodomain looks like a butterfly in 2D classification and low-resolution model. However, the monomer seems like a revolver with a solid center and two flexible ends as reported^35^. Model comparation showed that two subunits in homodimer are conformationally different from each other, furthermore only one of them adopts similar conformation as the monomer. Unfortunately, we could not solve the high resolution cryo-EM structure of homodimer for deeper analysis mostly due to the high flexibility of one subunit relative to the other one. Even though it has been proposed that dimerization is conserved in DBL domain receptor engagement for another DBL domain protein (RII-PvDBP) in *Plasmodium vivax*^42^. Whether the homodimer of VAR2CSA is physiologically relevant or just artificial due to overexpression needs to be further addressed.

Although the map density of the wing region can be clearly visualized in both our apo and complex structures, the low resolution and poor map quality due to the high flexibility make it hard to be interpreted. Compared with the map of the wing region in VAR2CSA ectodomain, the counterpart of the complex has been significantly improved, which makes it feasible for the flexible fitting of both DBL5ε and DBL6ε. According to our current biochemistry and cell biology experiments, there is no clear relevance observed between the wing region and the recognition of CSA by VAR2CSA at this stage. The conformational diversity adopted by the wing region could be speculated based on the 3D classification, which has shown multiple orientations relative to the core region (Fig. S3). However, the high flexibility of the wing region derived from the truncated VAR2CSA might be an artifact, because in terms of the full length VAR2CSA *in vivo*, DBL5ε and DBL6ε are physically connected to the transmembrane domain, which might restrict their movement to reduce their flexibility.

Over the past two decades since VAR2CSA was identified as a key virulent factor in PAM, the functional mechanism of VAR2CSA remains elusive due to its huge size and complexity until this year. Recently, two related works have been published in Nature Microbiology (NM) and Nature Communication (NC). Briefly, the cryo-EM structures of VAR2CSA ectodomain from *P. falciparum* strain FCR3 and VAR2CSA-CSA from *P. falciparum* strain NF54 were reported with CSA observed in NM paper. Meanwhile, the cryo-EM structures of VAR2CSA ectodomain from *P. falciparum* strain FCR3 and its complex with placental chondroitin sulfate were reported with no obvious ligand density in NC paper. Uniquely, our structural study showed that CSA binding induced significant conformational change to secure the ligand binding as described above, which was not reported in both papers. As for NM paper, one possible explanation is that they might catch a different conformation of the complex from ours. Another possibility is that VAR2CSA proteins in apo structure and complex structure are from two different *P. falciparum* strains with a sequence identity of 79%. As for NC paper, FCR3 VAR2CSA used share a sequence identity of 79% with 3D7 VAR2CSA used in our study, while no ligand density was reported in their complex structure. Most impressively, our work provided solid biological proofs to verify that 9 key residues in DBL2X were mainly responsible for CSA binding and cytoadherence.

In summary, the architecture and detailed molecular mechanism of cytoadherence to placenta or tumor cells through VAR2CSA were revealed using multi-disciplinary methods, which may facilitate PAM vaccine design and improve the delivery systems targeting both placenta and tumors.

## Methods

### VAR2CSA fragments expression and purification

VAR2CSA ectodomain coding sequence (residues 1-2630 from *P.falciparum* 3D7, GenBank ID: 811060) was synthesized with optimized codons for mammalian cell, and inserted into gp67-438-B vector with a N-terminal 6xHis Tag to prepare the bacmid and following recombinant baculovirus. The gp67-438-B modified from 438-B (Addgene) was used to express recombinant secretory proteins.

Sf9 cells were cultured in serum-free medium (SF900 II, Thermo Fisher) at 27°C with a speed of 130 rpm in a shaker (Yonglian). One liter of Sf9 cells (2 × 10^6^/mL confluence) was infected with 5 mL P3 baculoviral stock obtained following the manufacturer’s instructions (Invitrogen). The medium containing secreted protein was collected on the 4^th^ day (about 72-hour post-infection) and centrifuged for 20 min at 6,700 g to remove cell pellets. The supernatant was precipitated by saturated ammonium sulfate. And the precipitate was acquired by centrifugation for 40 min at 12,000 g, and resuspended in buffer A (20mM MES, pH6.5). The sample was mixed with pre-equilibrated Ni-NTA Agarose beads (GE Healthcare) at a ratio of 2 mL beads per liter medium and stirred for 2 hours at 4°C. The slurry was loaded onto a 15-mL gravity column (Bio-Rad) and washed with wash buffer (20 mM MES pH 6.5, 500 mM NaCl, and 5% v/v glycerol) for ~30 column volumes. The beads were further washed using wash buffer supplemented with 10 mM imidazole until no trace of protein was detected in the flowthrough. VAR2CSA was eluted by wash buffer supplemented with 200 mM imidazole. It was then concentrated (Amicon Ultra-15 30,000 MWCO, Millipore) to ~500 μL and loaded onto a Superose6 increase 10/300 GL (GE Healthcare) equilibrated with buffer B (20 mM MES pH 6.5, 250 mM NaCl). The peak fractions containing final pure protein were collected, pooled, and concentrated to 3 mg/mL for the following experiments.

VAR2CSA-50-962 and VAR2CSA-550-962 were amplified and subcloned into the pET20b vector between NdeI and XhoI restriction sites with a C-terminal 6xHis tag. VAR2CSA-50-962-9-site-mutation was generated via the substitution of the synthesized fragment with 9 key residues mutated (N557D, K561E, K562E, N576D, K828E, R829E, Q832E, K835E, R846E). The above proteins were expressed in Rosetta-gamiB (DE3) of *E. coli*. (WEIDI Biotechnology). Briefly, when the recombinant bacteria grow to the OD value around 0.6 at 37°C, the temperature was lowered to 16°C for about half an hour. IPTG was then added to the culture at a final concentration of 0.2 mM. Cells were collected after 20-hour incubation. Cell pellet were resuspended in wash buffer and lysed using Emulsiflex homogeniser (YongLian). The proteins were purified by sequential chromatography using Ni-NTA affinity, HiTrap S, gel filtration columns (GE Healthcare). According to the separation efficiency, Superose6 increase 10/300 GL was used for VAR2CSA-50-962 and VAR2CSA-50-962-9-site-mutation, and Superdex 75 10/300 GL for VAR2CSA-550-962.

CSA extracted from bovine trachea (Sigma, C9819) was used as the ligand for VAR2CSA. Various VAR2CSA proteins were incubated with CSA solution (10 mg/mL) overnight on ice and further separated by Superose6 increase 10/300 GL. The complex fractions were collected, pooled and concentrated to 3 mg/mL for the following experiments.

### Cryo-samples preparation of VAR2CSA ectodomain

The VAR2CSA ectodomain was cross-linked and purified using Grafix^43^. Briefly, the glycerol gradient was prepared using light buffer (20 mM MES pH 6.5, 250 mM NaCl, 15% (v/v) glycerol) and heavy buffer (20 mM MES pH 6.5, 250 mM NaCl, 0.05% glutaraldehyde, 35% (v/v) glycerol). The samples were centrifuged at 38,000 rpm for 16 hours at 4°C using a Beckman SW41 Ti rotor. Subsequently, 200 μL per fraction were harvested and the cross-linking reaction was terminated by adding quench buffer (1 M Tris pH 6.5, 250 mM NaCl) to a final concentration of 40 mM Tris. Fractions containing crosslinked VAR2CSA monomer were pooled, concentrated, and dialyzed in buffer B. The cross-linked VAR2CSA monomer about 0.4 mg/mL was applied to the preparation of cryo-EM grids.

For negative staining EM analysis, carbon-coated copper grids were processed using a PELCO easiGlow (TED PELLA) cleaning system with a power of 30 W and a plasma current of 30 mA in the air for 30 s. Subsequently, samples (8 μL at a concentration of ~0.02 mg/mL) were applied onto the grids above and stained twice using 2% (w/v) uranyl acetate solution at room temperature for the following examination via 200kV TEM (TF20, FEI).

As for the preparation of cryo-EM grids, Amorphous Alloy Film Au300 R1.2/1.3 grids (CryoMatrix) were processed in the H_2_/O_2_ mixture for 30 s using a Gatan 950 Solarus plasma cleaning system with a power of 5 W. Then cross-linked VAR2CSA ectodomain (3 μL at a concentration of ~0.4 mg/mL) was applied to the grids for instant incubation under a relative humidity of 100% at 4 °C. Next, the grids were blotted for 2 s with a blot force of 1 in a Vitrobot Mark IV (Thermo Fisher) and plunge-frozen in liquid ethane. Similarly, VAR2CSA-CSA (3 μL at a concentration of ~0.5 mg/mL) was blotted for 3 s with a blot force of −2 and plunge-frozen in liquid ethane.

### Cryo-EM data collection and Image processing

The cryo-EM grids of cross-linked VAR2CSA ectodomain and VAR2CSA-CSA were firstly evaluated using a 200kV Talos Arctica microscope (Thermo Fisher). Cryo-EM datasets were collected on 300 kV Titan Krios microscope (Thermo Fisher) equipped with a Gatan K2 Summit direct electron detector and a 20-eV slit GIF Quantum energy filter (Gatan). The cryo-EM images were automatically recorded in the super-resolution counting mode using Serial-EM^44^ software with a nominal magnification of 130,000 x (a super-resolution pixel size of 0.522 Å), and with a defocus ranging from −1.2 to −2.2 μm. Each micrograph stack was dose-fractionated into 36 frames with a total electron dose of ~ 57.6 e^-^/ Å^2^ and a total exposure time of 7.2 s. Drift and beam-induced motion of the super-resolution movie stacks were corrected using MotionCor2^45^ and binned twofold to a calibrated pixel size of 1.044 Å/pix, with both the dose weighted and non-dose weighted micrographs saved at the same time. The defocus values were estimated by Gctf^46^ using the non-dose weighted micrographs. Other procedures of cryo-EM data processing were performed using RELION v3.0^47^.

A total of 1,965 movies were recorded for the cross-linked VAR2CSA ectodomain. Among them, 1,800 micrographs were selected for further processing due to appropriate range of rlnDefocusU (5000-30000) and rlnCtfMaxResolution (2-6). Selected 2D class averages from 200kV cryo-EM data were low-pass filtered to 20 Å and used as templates for autopicking. Then an initial set of 984,517 particles were extracted for 2D classification. After several rounds of 2D classification, the relatively good classes with 960,039 particles were selected for three-dimensional reconstruction. Initially, one major class (~62% particles) was separated after 3D classification. To further improve the resolution, a second round of 3D classification with mask was carried out with 304,160 particles selected for 3D autorefinement and postprocessing (the B-factor automatically estimated). The final resolution was evaluated according the gold-standard Fourier shell correlation (threshold = 0.143)^48^. Although the overall resolution is up to 3.6 Å, the density of the wing region (DBL5ε and DBL6ε) is quite weak due to high flexibility. The local refinement strategy for focused classification and refinement was carried out for the whole structure divided into two half parts. Consequently, the resolution of the core region could be improved to 3.1 Å by merging two local-refined maps. However, the map quality of the wing region remained unimproved, which made it hardly interpreted by *de novo* model building. The local resolution was evaluated by ResMap^49^.

For VAR2CSA-CSA, 3,058 out of 3,356 micrographs were selected for further processing using similar procedures as above. Briefly, 1,163,625 auto-picked particles were extracted for 2D classification. After several rounds of 2D and 3D classification, 241,954 particles were selected for 3D auto-refinement and postprocessing. The final map was reconstructed at a resolution of 3.4 Å. Compared with apo structure, the density of the wing region has been significantly improved, which makes it feasible for the flexible fitting of both DBL5ε and DBL6ε. The resolution of the core region could be improved to 3.1 Å by merging local-refined maps.

### Model building and refinement

The density map of VAR2CSA ectodomain was firstly interpreted using Phenix.map_to_model^50^ to generate the initial model with the most helixes, which has been further improved by manual model building using COOT^51^. The structure of VAR2CSA-CSA was determined using the apo structure as a reference and manual model building in COOT with various refinement strategies. The structures have been further validated by validation module in Phenix (Table S2). All structural figures were prepared using UCSF Chimera^52^, UCSF ChimeraX^53^ or PyMOL^54^.

### Evaluation of protein-protein interaction by Octet RED 96

CSA were biotinylated and loaded onto SA sensor (Pall corporation). VAR2CSA fragments and mutant were then added for real-time association and dissociation analysis using Octet RED 96 (Fortebio) at room temperature. Data Analysis Octet was used for data processing.

### Confocal fluorescence microscopy

Cells were pre-seeded on a slide, washed with PBS and fixed using 4% PFA/PBS on ice. After fixation, cells were blocked in 2.5% FBS for 30 min on ice and incubated with recombinant VAR2CSA proteins diluted in PBS with 0.25% FBS for 1 hour on ice. After washing with PBS by three times, the specimens were incubated with anti-His-FITC antibody (1:500) for 1 hour at room temperature in the dark and washed as described before. DAPI was applied to stain and locate the cell nuclei. Slides were further analyzed using a confocal microscope. Negative controls (Mock) were prepared with the same procedures except the incubation of recombinant VAR2CSA fragments.

## Acknowledgements

We would like to pay an exceptional tribute to the staff members of the Cryo-EM facility of Fudan University (Fenglin Campus) and the Bio-Electron Microscopy Facility of ShanghaiTech University for their generous support in cryo-sample preparation and data collection. We thank the staffs, particularly Jie Li and Lei Zhang, of the Large-scale Protein Preparation System at the National Facility for Protein Science in Shanghai (NFPS), Zhangjiang Lab, China for providing technical support and assistance in data collection and analysis using Octet RED 96. We also thank the scientific core facility members in Institut Pasteur of Shanghai, CAS for their help in negative staining TEM and confocal fluorescence microscopy. This work was funded by the National Key R&D Program of China (2018YFA0507303; 2017YFC0840300); the Strategic Priority Research Program of CAS (XDB29010205); and the National Natural Science Foundation of China (NSFC) (31770816; 82025023; 31771455; and 31970146).

## Author contributions

L. W., Z. C. and L. J. conceived, designed, and coordinated this project; W. W.^**1**^, J. Z., Q. J., and L. C. expressed and purified the N-terminal domain; W. W.^**1**^ and Q. X. prepared the rest samples containing ectodomain, complex, and mutations; W. W.^**1**^, J. S., and Z. W. carried out the Octet RED 96 and confocal fluorescence microscopy experiments; Z. C., L. S., X. Y., Z. W., Y. G., X. Z., and W. W. ^**3**^ prepared the cryo-samples and collected cryo-EM data; Z. C., Z. W., Y. G., Y. T., A. D., and W. W.^**3**^ processed cryo-EM data; L. W., and Z. W. built and refined the structure models; the manuscript was written by L. W., W. W.^**1**^, Z. W., L. S., Z. C., X. Y., Y. G.; All authors discussed the experiments and results, read, and approved the manuscript.

## Competing interests

All the authors declare no conflict of interest.

## Materials & Correspondence

Correspondence and requests for materials should be addressed to L. J., Z. C. or L. W.

## Data availability

The atomic coordinates and electron microscopy data have been deposited in the RCSB Protein Data Bank and Electron Microscopy Data Bank under the following entries: 3D7 VAR2CSA ectodomain (PDB-ID 7FAS, EMD-31505), 3D7 VAR2CSA-CSA complex (PDB-ID 7FAP, EMD-31504).

## Notes

### Competing Interest Statement

The authors have declared no competing interest.

## Reference

1 Guerra, C. A. et al. The limits and intensity of Plasmodium falciparum transmission: implications for malaria control and elimination worldwide. PLoS Med 5, e38, doi:10.1371/journal.pmed.0050038 (2008).

2 Brabin, B. J., Hakimi, M. & Pelletier, D. An analysis of anemia and pregnancy-related maternal mortality. J Nutr 131, 604S–614S; discussion 614S-615S, doi:10.1093/jn/131.2.604S (2001).

3 Huynh, B. T. et al. Influence of the timing of malaria infection during pregnancy on birth weight and on maternal anemia in Benin. Am J Trop Med Hyg 85, 214–220, doi:10.4269/ajtmh.2011.11-0103 (2011).

4 World Health, O. World malaria report 2019. xxxix, 185 p. (World Health Organization, 2019).

5 Salanti, A. et al. Selective upregulation of a single distinctly structured var gene in chondroitin sulphate A-adhering Plasmodium falciparum involved in pregnancy-associated malaria. Mol Microbiol 49, 179–191, doi:10.1046/j.1365-2958.2003.03570.x (2003).

6 Elliott, S. R. et al. Cross-reactive surface epitopes on chondroitin sulfate A-adherent Plasmodium falciparum-infected erythrocytes are associated with transcription of var2csa. Infect Immun 73, 2848–2856, doi:10.1128/IAI.73.5.2848-2856.2005 (2005).

7 Miller, L. H., Baruch, D. I., Marsh, K. & Doumbo, O. K. The pathogenic basis of malaria. Nature 415, 673–679, doi:10.1038/415673a (2002).

8 Viebig, N. K. et al. A single member of the Plasmodium falciparum var multigene family determines cytoadhesion to the placental receptor chondroitin sulphate A. EMBO Rep 6, 775–781, doi:10.1038/sj.embor.7400466 (2005).

9 Fried, M. & Duffy, P. E. Designing a VAR2CSA-based vaccine to prevent placental malaria. Vaccine 33, 7483–7488, doi:10.1016/j.vaccine.2015.10.011 (2015).

10 Alkhalil, A., Achur, R. N., Valiyaveettil, M., Ockenhouse, C. F. & Gowda, D. C. Structural requirements for the adherence of Plasmodium falciparum-infected erythrocytes to chondroitin sulfate proteoglycans of human placenta. J Biol Chem 275, 40357–40364, doi:10.1074/jbc.M006399200 (2000).

11 Goel, S. & Gowda, D. C. How specific is Plasmodium falciparum adherence to chondroitin 4-sulfate? Trends Parasitol 27, 375–381, doi:10.1016/j.pt.2011.03.005 (2011).

12 Salanti, A. et al. Targeting Human Cancer by a Glycosaminoglycan Binding Malaria Protein. Cancer Cell 28, 500–514, doi:10.1016/j.ccell.2015.09.003 (2015).

13 Srivastava, A., Durocher, Y. & Gamain, B. Expressing full-length functional PfEMP1 proteins in the HEK293 expression system. Methods Mol Biol 923, 307–319, doi:10.1007/978-1-62703-026-7_22 (2013).

14 Srivastava, A. et al. Full-length extracellular region of the var2CSA variant of PfEMP1 is required for specific, high-affinity binding to CSA. Proc Natl Acad Sci U S A 107, 4884–4889, doi:10.1073/pnas.1000951107 (2010).

15 Khunrae, P. et al. Full-length recombinant Plasmodium falciparum VAR2CSA binds specifically to CSPG and induces potent parasite adhesion-blocking antibodies. J Mol Biol 397, 826–834, doi:10.1016/j.jmb.2010.01.040 (2010).

16 Dechavanne, S. et al. Parity-dependent recognition of DBL1X-3X suggests an important role of the VAR2CSA high-affinity CSA-binding region in the development of the humoral response against placental malaria. Infect Immun 83, 2466–2474, doi:10.1128/IAI.03116-14 (2015).

17 Avril, M. et al. Characterization of anti-var2CSA-PfEMP1 cytoadhesion inhibitory mouse monoclonal antibodies. Microbes Infect 8, 2863–2871, doi:10.1016/j.micinf.2006.09.005 (2006).

18 Nielsen, M. A. et al. Induction of adhesion-inhibitory antibodies against placental Plasmodium falciparum parasites by using single domains of VAR2CSA. Infect Immun 77, 2482–2487, doi:10.1128/IAI.00159-09 (2009).

19 Avril, M. et al. Immunization with VAR2CSA-DBL5 recombinant protein elicits broadly cross-reactive antibodies to placental Plasmodium falciparum-infected erythrocytes. Infect Immun 78, 2248–2256, doi:10.1128/IAI.00410-09 (2010).

20 Badaut, C. et al. Towards the rational design of a candidate vaccine against pregnancy associated malaria: conserved sequences of the DBL6epsilon domain of VAR2CSA. PLoS One 5, e11276, doi:10.1371/journal.pone.0011276 (2010).

21 Avril, M., Cartwright, M. M., Hathaway, M. J. & Smith, J. D. Induction of strain-transcendent antibodies to placental-type isolates with VAR2CSA DBL3 or DBL5 recombinant proteins. Malar J 10, 36, doi:10.1186/1475-2875-10-36 (2011).

22 Doritchamou, J. Y. et al. VAR2CSA Domain-Specific Analysis of Naturally Acquired Functional Antibodies to Plasmodium falciparum Placental Malaria. J Infect Dis 214, 577–586, doi:10.1093/infdis/jiw197 (2016).

23 Sirima, S. B. et al. PRIMVAC vaccine adjuvanted with Alhydrogel or GLA-SE to prevent placental malaria: a first-in-human, randomised, double-blind, placebo-controlled study. Lancet Infect Dis 20, 585–597, doi:10.1016/S1473-3099(19)30739-X (2020).

24 Mordmuller, B. et al. First-in-human, Randomized, Double-blind Clinical Trial of Differentially Adjuvanted PAMVAC, A Vaccine Candidate to Prevent Pregnancy-associated Malaria. Clin Infect Dis 69, 1509–1516, doi:10.1093/cid/ciy1140 (2019).

25 Zhang, B. et al. Placenta-specific drug delivery by trophoblast-targeted nanoparticles in mice. Theranostics 8, 2765–2781, doi:10.7150/thno.22904 (2018).

26 Zhao, K. et al. Targeting delivery of partial VAR2CSA peptide guided N-2-Hydroxypropyl trimethyl ammonium chloride chitosan nanoparticles for multiple cancer types. Mater Sci Eng C Mater Biol Appl 106, 110–171, doi:10.1016/j.msec.2019.110171 (2020).

27 Bang-Christensen, S. R. et al. Capture and Detection of Circulating Glioma Cells Using the Recombinant VAR2CSA Malaria Protein. Cells 8, doi:10.3390/cells8090998 (2019).

28 Agerbaek, M. O. et al. The VAR2CSA malaria protein efficiently retrieves circulating tumor cells in an EpCAM-independent manner. Nat Commun 9, 3279, doi:10.1038/s41467-018-05793-2 (2018).

29 Hsieh, F. L. et al. The structural basis for CD36 binding by the malaria parasite. Nat Commun 7, 12837, doi:10.1038/ncomms12837 (2016).

30 Singh, K. et al. Structure of the DBL3x domain of pregnancy-associated malaria protein VAR2CSA complexed with chondroitin sulfate A. Nat Struct Mol Biol 15, 932–938, doi:10.1038/nsmb.1479 (2008).

31 Higgins, M. K. The structure of a chondroitin sulfate-binding domain important in placental malaria. J Biol Chem 283, 21842–21846, doi:10.1074/jbc.C800086200 (2008).

32 Gangnard, S. et al. Structure of the DBL3X-DBL4epsilon region of the VAR2CSA placental malaria vaccine candidate: insight into DBL domain interactions. Sci Rep 5, 14868, doi:10.1038/srep14868 (2015).

33 Khunrae, P., Philip, J. M., Bull, D. R. & Higgins, M. K. Structural comparison of two CSPG-binding DBL domains from the VAR2CSA protein important in malaria during pregnancy. J Mol Biol 393, 202–213, doi:10.1016/j.jmb.2009.08.027 (2009).

34 Gangnard, S. et al. Structural and immunological correlations between the variable blocks of the VAR2CSA domain DBL6epsilon from two Plasmodium falciparum parasite lines. J Mol Biol 425, 1697–1711, doi:10.1016/j.jmb.2013.02.014 (2013).

35 Bewley, M. C., Gautam, L., Jagadeeshaprasad, M. G., Gowda, D. C. & Flanagan, J. M. Molecular architecture and domain arrangement of the placental malaria protein VAR2CSA suggests a model for carbohydrate binding. J Biol Chem, doi:10.1074/jbc.RA120.014676 (2020).

36 Terwilliger, T. C., Adams, P. D., Afonine, P. V. & Sobolev, O. V. Cryo-EM map interpretation and protein model-building using iterative map segmentation. Protein Sci 29, 87–99, doi:10.1002/pro.3740 (2020).

37 Muthusamy, A. et al. Structural characterization of the bovine tracheal chondroitin sulfate chains and binding of Plasmodium falciparum-infected erythrocytes. Glycobiology 14, 635–645, doi:10.1093/glycob/cwh077 (2004).

38 Clausen, T. M. et al. Structural and functional insight into how the Plasmodium falciparum VAR2CSA protein mediates binding to chondroitin sulfate A in placental malaria. J Biol Chem 287, 23332–23345, doi:10.1074/jbc.M112.348839 (2012).

39 Koshland, D. E. Application of a Theory of Enzyme Specificity to Protein Synthesis. Proc Natl Acad Sci U S A 44, 98–104, doi:10.1073/pnas.44.2.98 (1958).

40 Dahlback, M. et al. The chondroitin sulfate A-binding site of the VAR2CSA protein involves multiple N-terminal domains. J Biol Chem 286, 15908–15917, doi:10.1074/jbc.M110.191510 (2011).

41 Bigey, P. et al. The NTS-DBL2X region of VAR2CSA induces cross-reactive antibodies that inhibit adhesion of several Plasmodium falciparum isolates to chondroitin sulfate A. J Infect Dis 204, 1125–1133, doi:10.1093/infdis/jir499 (2011).

42 Batchelor, J. D., Zahm, J. A. & Tolia, N. H. Dimerization of Plasmodium vivax DBP is induced upon receptor binding and drives recognition of DARC. Nat Struct Mol Biol 18, 908–914, doi:10.1038/nsmb.2088 (2011).

43 Kastner, B. et al. GraFix: sample preparation for single-particle electron cryomicroscopy. Nature Methods 5, 53–55, doi:10.1038/nmeth1139 (2008).

44 Mastronarde, D. N. Automated electron microscope tomography using robust prediction of specimen movements. Journal of Structural Biology 152, 36–51, doi:https://doi.org/10.1016/j.jsb.2005.07.007 (2005).

45 Zheng, S. Q. et al. MotionCor2: anisotropic correction of beam-induced motion for improved cryo-electron microscopy. Nature Methods 14, 331–332, doi:10.1038/nmeth.4193 (2017).

46 Zhang, K. Gctf: Real-time CTF determination and correction. Journal of Structural Biology 193, 1–12, doi:https://doi.org/10.1016/j.jsb.2015.11.003 (2016).

47 Scheres, S. H. W. RELION: Implementation of a Bayesian approach to cryo-EM structure determination. Journal of Structural Biology 180, 519–530, doi:https://doi.org/10.1016/j.jsb.2012.09.006 (2012).

48 Scheres, S. H. W. & Chen, S. Prevention of overfitting in cryo-EM structure determination. Nature Methods 9, 853–854, doi:10.1038/nmeth.2115 (2012).

49 Kucukelbir, A., Sigworth, F. J. & Tagare, H. D. Quantifying the local resolution of cryo-EM density maps. Nature Methods 11, 63–65, doi:10.1038/nmeth.2727 (2014).

50 Afonine, P. V. et al. Towards automated crystallographic structure refinement with phenix.refine. Acta Crystallogr D Biol Crystallogr 68, 352–367, doi:10.1107/S0907444912001308 (2012).

51 Brown, A. et al. Tools for macromolecular model building and refinement into electron cryo-microscopy reconstructions. Acta Crystallogr D Biol Crystallogr 71, 136–153, doi:10.1107/S1399004714021683 (2015).

52 Pettersen, E. F. et al. UCSF Chimera—A visualization system for exploratory research and analysis. Journal of Computational Chemistry 25, 1605–1612, doi:https://doi.org/10.1002/jcc.20084 (2004).

53 Goddard, T. D. et al. UCSF ChimeraX: Meeting modern challenges in visualization and analysis. Protein Science 27, 14–25, doi:https://doi.org/10.1002/pro.3235 (2018).

54 Schrodinger, LLC. The PyMOL Molecular Graphics System, Version 1.8 (2015).

