## Supplementary information for "The molecular mechanism of cytoadherence to placenta or tumor cells through VAR2CSA from *Plasmodium falciparum*"

### Supplementary materials

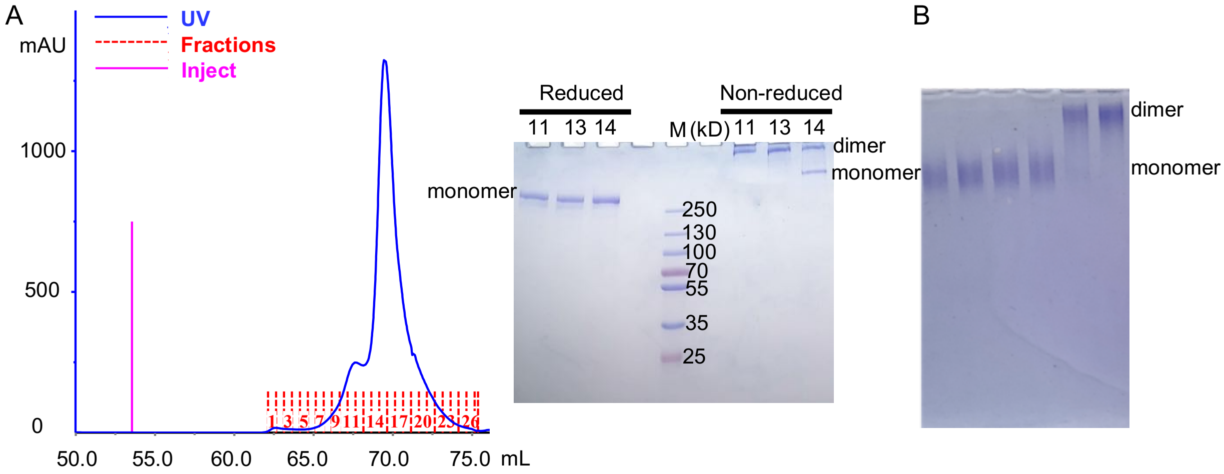


**Fig. S1 The dimerization of VAR2CSA ectodomain. A**, Chromatogram of VAR2CSA ectodomain in Superose6 increase 10/300 column. Fractions were further evaluated using reduced and non-reduced SDS-PAGE. **B**, Crosslinked VAR2CSA ectodomain using Grafix showed the presence of both monomer and dimer.

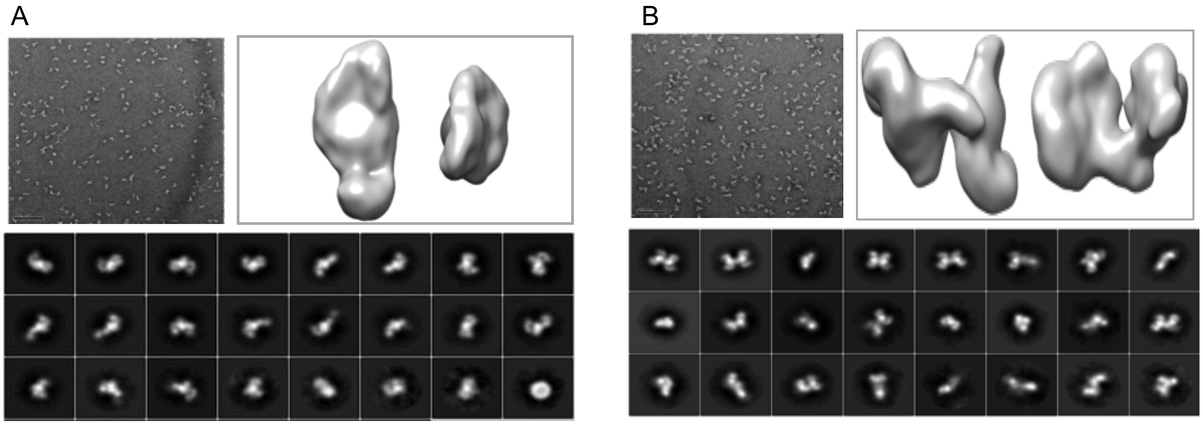


**Fig. S2 Negative staining model of VAR2CSA ectodomain in monomer (A) or dimer (B).** Up left, imaging picture; Down, 2D classification; up right, low resolution model.

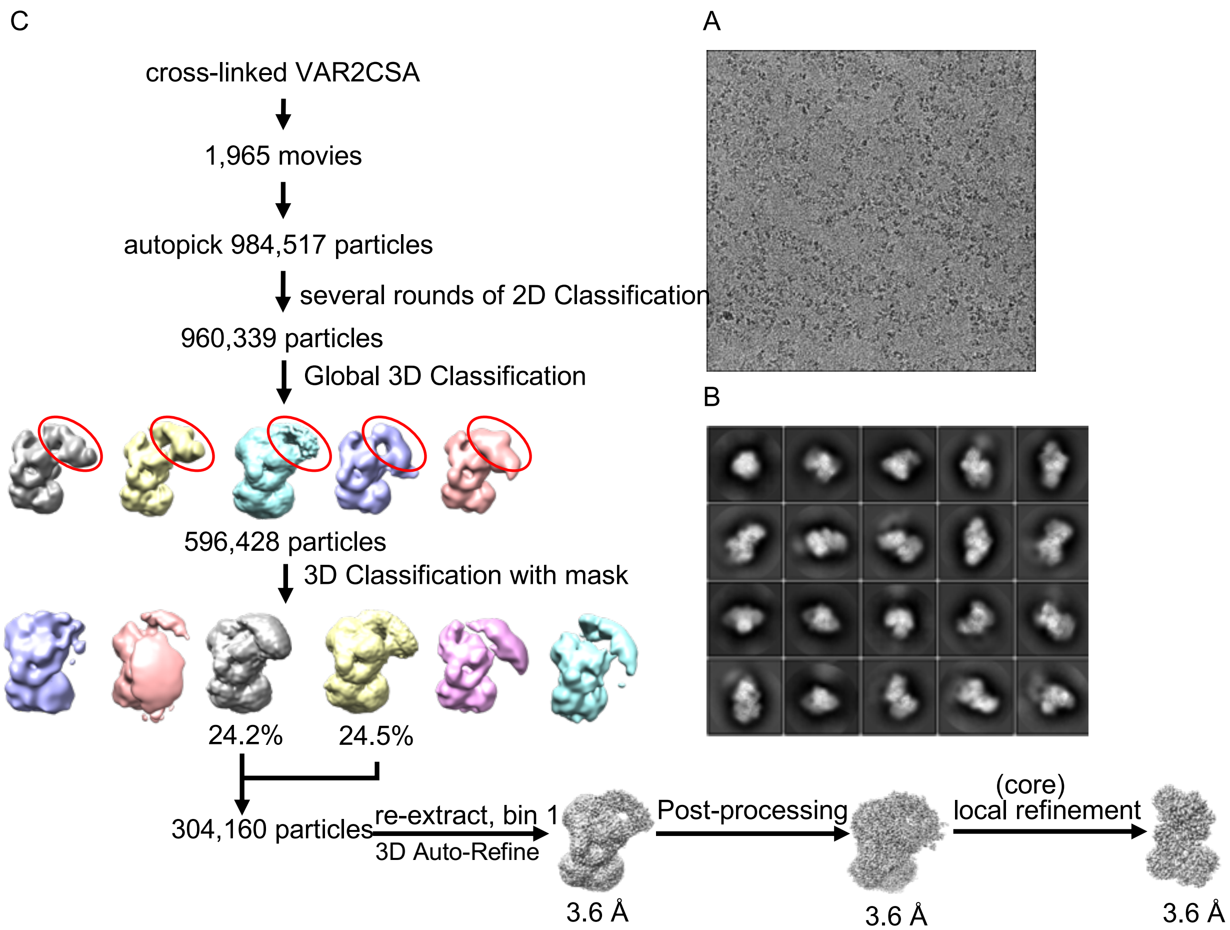


**Fig. S3 Data collection and image processing of VAR2CSA ectodomain.** **A**, Representative images of cryo-EM dataset. **B**, 2D classification result. **C**, The flowchart of the cryo-EM image processing and 3D reconstruction for cross-linked VAR2CSA ectodomain. The highly flexible wing region highlighted in red oval in global 3D classification.

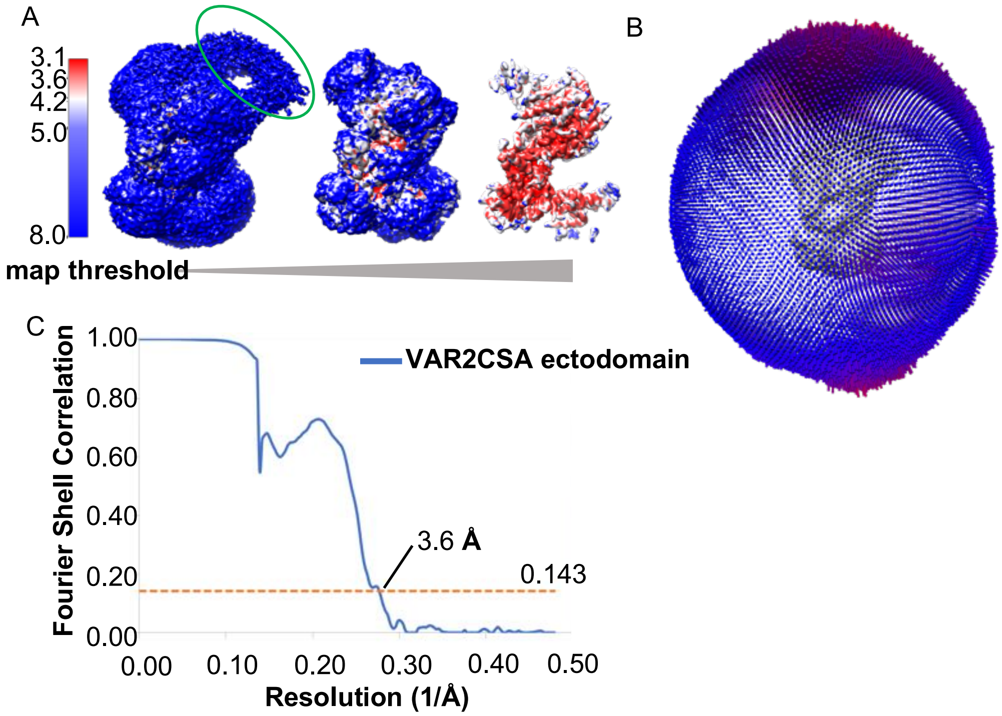


**Fig. S4 Cryo-EM analysis of VAR2CSA ectodomain. A**, Local resolution of VAR2CSA ectodomain under different threshold of cryo-EM density map estimated by ResMap. Flexible wing region highlighted in red oval. **B**, Angular distribution of cryo-EM samples of VAR2CSA ectodomain. **C**, Gold standard-FSC curve of the refined map.

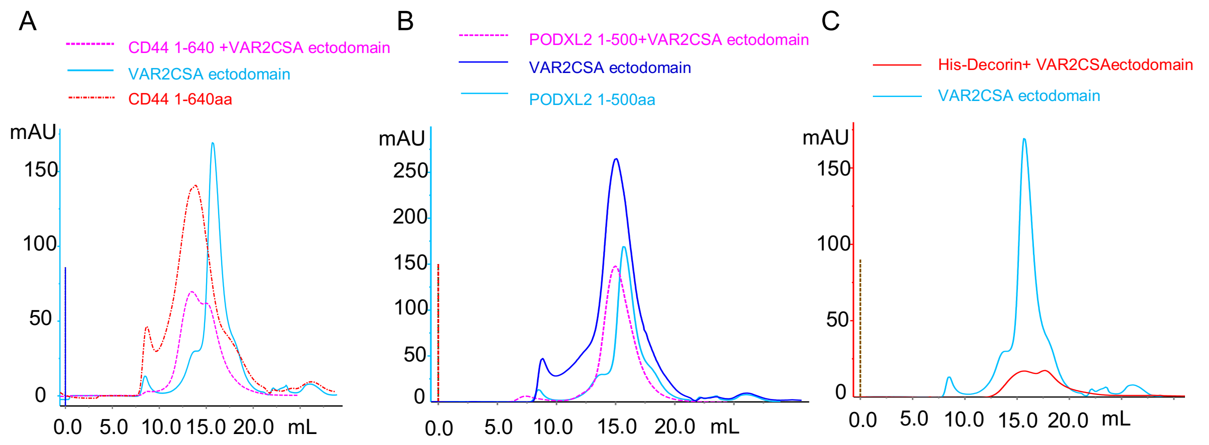


**Fig.S5 The interaction between potential glycoproteins and VAR2CSA ectodomain validated using gel filtration chromatography.** Chromatograms of VAR2CSA ectodomain in the presence or absence of CD44 (**A)**, PODXL2 (**B**), Decorin (**C**) using a Superose6 increase 10/300 GL column.

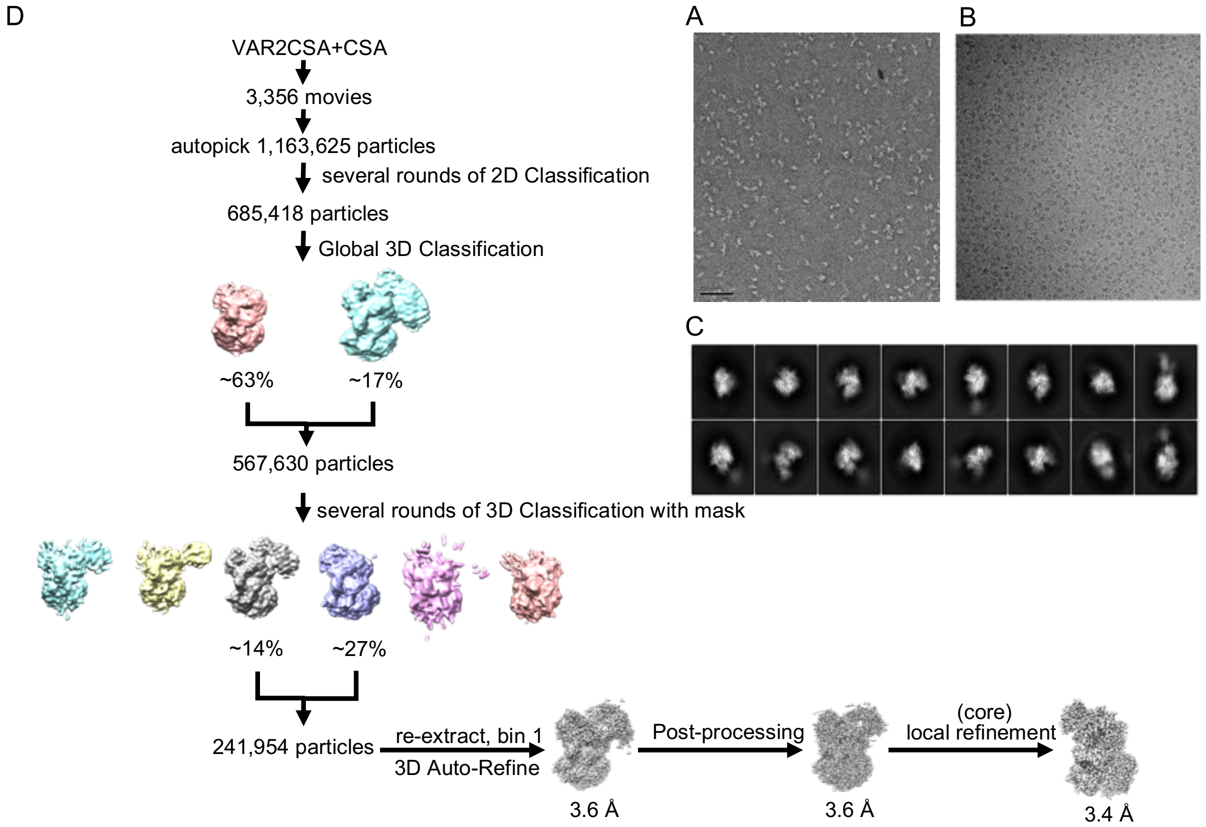


**Fig. S6 Data collection and image processing of VAR2CSA-CSA. (A-C)** Representative images of negative staining (**A**), cryo-EM dataset (**B**), and 2D classification (**C**). **D**, Flowchart of the cryo-EM image processing and 3D reconstruction for VAR2CSA-CSA.

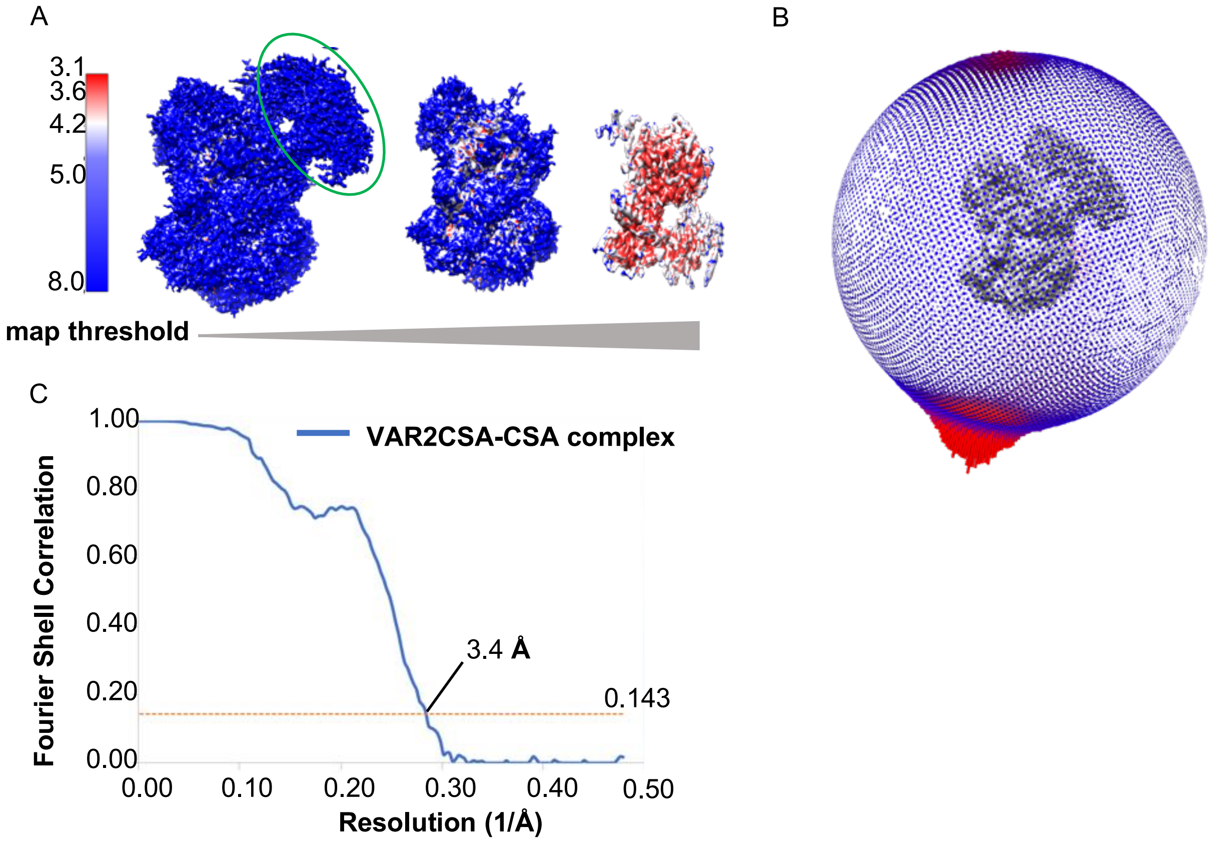


**Fig. S7 Cryo-EM analysis of VAR2CSA-CSA. A**, Local resolution of VAR2CSA-CSA under different threshold of cryo-EM density map estimated by ResMap. Flexible wing region highlighted in green oval. **B**, Angular distribution of cryo-EM samples of VAR2CSA-CSA. **C**, Gold standard-FSC curve of the refined map.

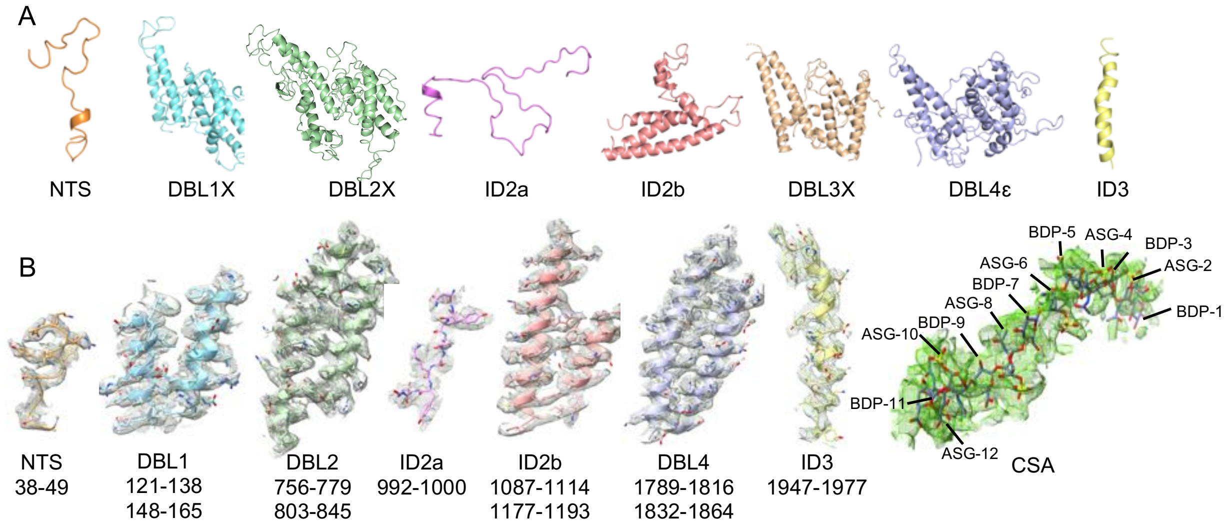


**Fig. S8 Model composition and map density validation for VAR2CSA-CSA. A**, Ribbon diagrams of individual domains in the structure of VAR2CSA-CSA core region. **B**, Representative cryo-EM density maps from VAR2CSA-CSA filled with structure models with side chains as sticks.

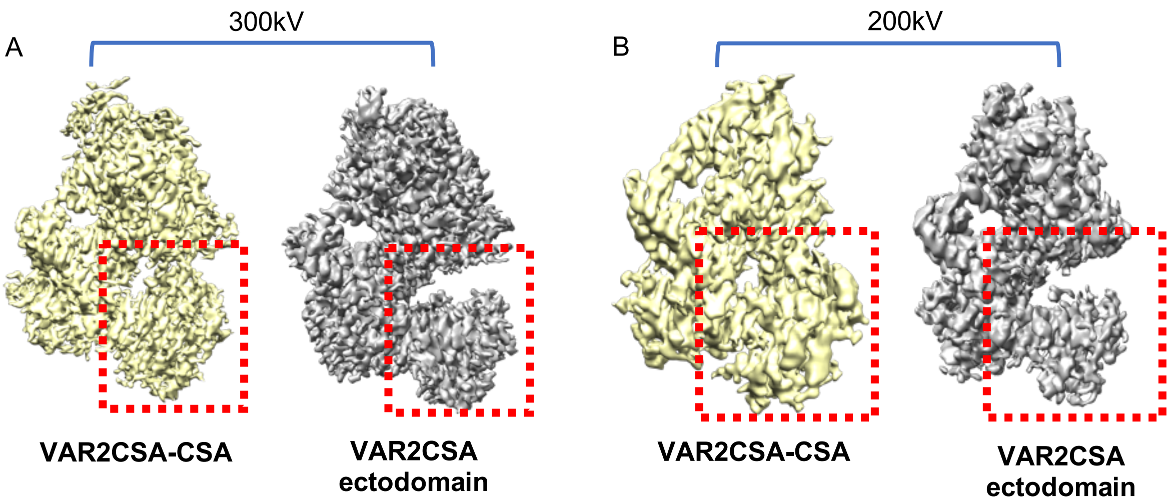


**Fig. S9 Cryo-EM density maps represent the conformational change of VAR2CSA ectodomain upon CSA binding.** Cryo-EM density maps reconstructed from 300 kV cryo-EM dataset (**A**) and 200 kV cryo-EM dataset (**B**) both displayed major differences highlighted within red dotted rectangles. VAR2CSA-CSA were colored in yellow and VAR2CSA ectodomain in grey.

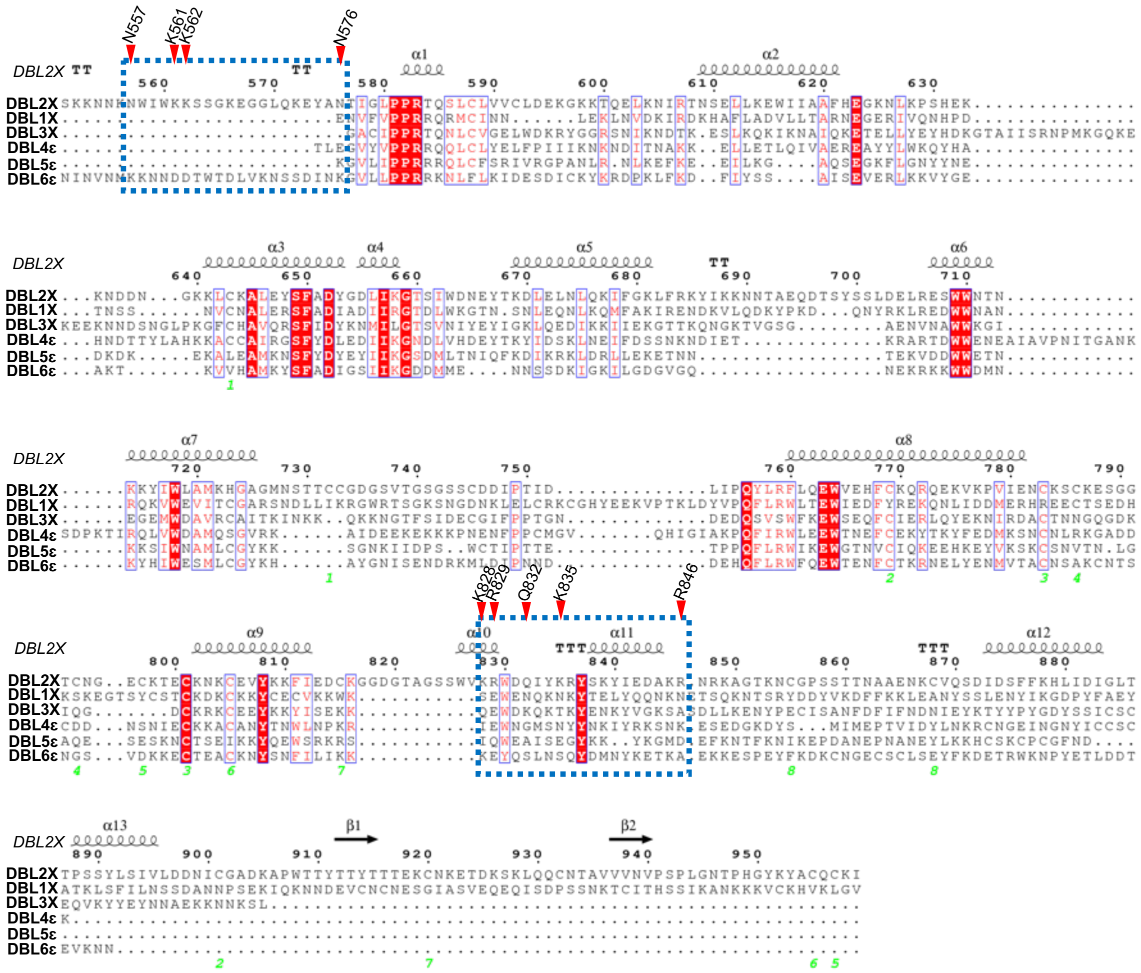

**Fig. S10 Sequence alignment of six individual VAR2CSA DBL domain.** The nine key residues of DBL2X involving in CSA binding were labeled and indicated by red triangles within the blue dotted rectangles. Secondary structures were labeled according to the structure of DBL2X using Espript3 online server.

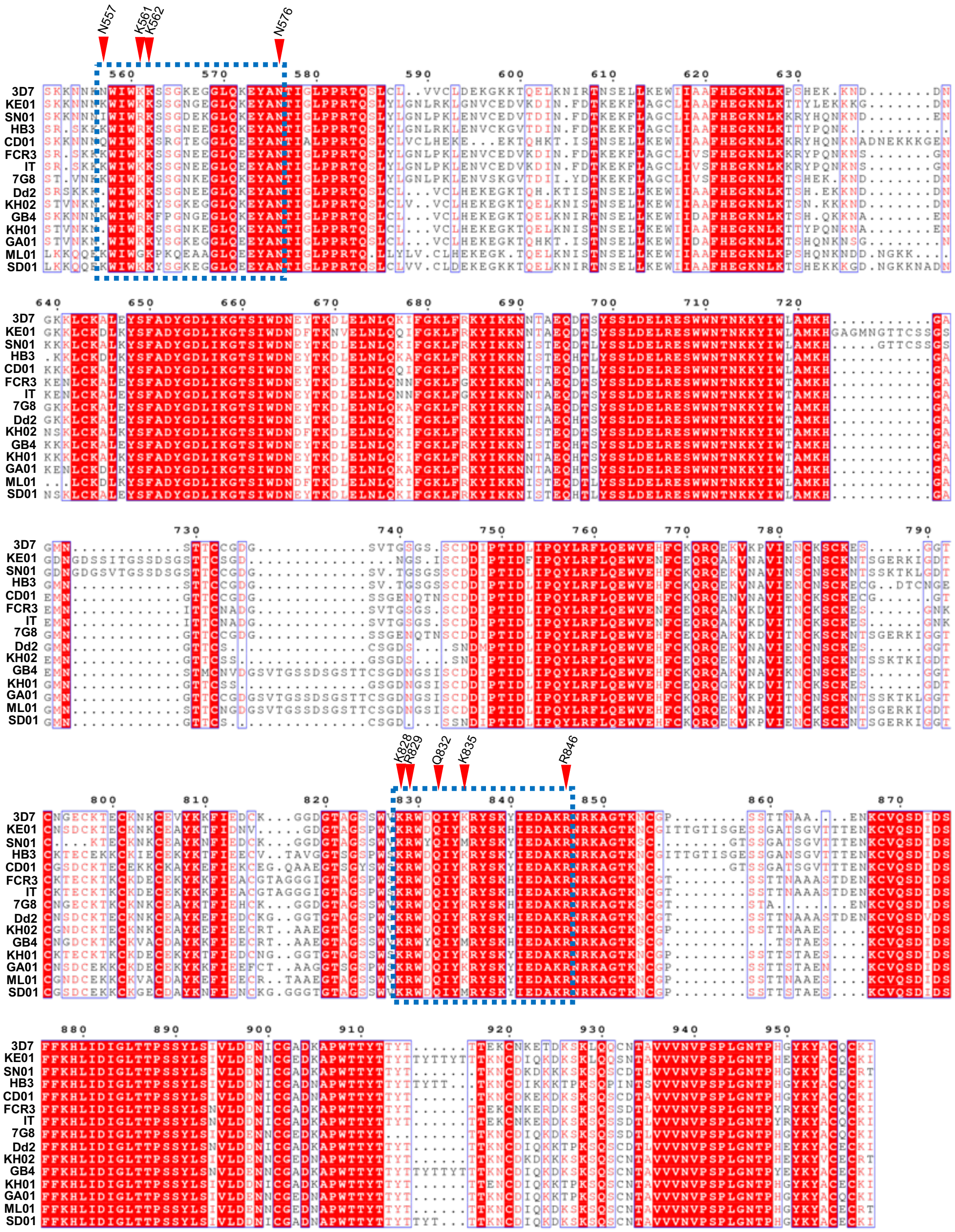


**Fig. S11 The sequence alignment of VAR2CSA DBL2X domains from various *Plasmodium falciparum* species.** 9-key-residue for binding CSA were highlighted.

**Table S1. The key residues in VAR2CSA ectodomain responsible for interacting with CSA.**

| Domain | Residue | Interaction unit within CSA |
|---|---|---|
| NTS | Y44 | ASG-2 |
| | Y45 | BDP-1 |
| | N49 | BDP-1/ASG-2 |
| DBL2X | N557 | ASG-12 |
| | K561 | ASG-10 |
| | K562 | ASG-8 |
| | N576 | ASG-8 |
| | K828 | ASG-8 |
| | R829 | BDP-9 |
| | Q832 | BDP-7 |
| | K835 | ASG-4 |
| | R846 | BDP-1 |
| DBL4ε | E1880 | ASG-4 |
| | K1889 | ASG-4/BDP-5 |
| | R1890 | BDP-3 |
| | Y1899 | BDP-1 |

**Table S2. Statistics of cryo-EM data collection and refinements**

| | VAR2CSA ectodomain | VAR2CSA-CSA |
|---|---|---|
| **Data collection and processing** | | |
| Magnification | 130,000x | 130,000x |
| Voltage (kV) | 300 | 300 |
| Electron exposure ($e^-$ /pix/s) | ~8.0 | ~8.0 |
| Exposure rate ($e^-$ /Å$^2$ ) | ~50 | ~50 |
| Number of frames per movie | 36 | 36 |
| Energy filter slit width (eV) | 20 | 20 |
| Automation software | SerialEM | SerialEM |
| Defocus range (µm) | -1.2~-2.2 | -1.2~-2.2 |
| Pixel size (Å) | 1.044 | 1.044 |
| Symmetry imposed | C1 | C1 |
| Micrographs (no.) | 1,965 | 3,356 |
| Total of extracted particles (no.) | 984,517 | 1,163,625 |
| Total of refined particles (no.) | 304,160 | 241,954 |
| Map resolution (Å) | | |
| FSC threshold | 0.143 | 0.143 |
| Map resolution range (Å) | 3.6 | 3.4 |
| **Refinement** | | |
| Map sharpening B-factor (Å$^2$ ) | -85.59 | -123.93 |
| Model composition | | |
| Nonhydrogen atoms | 10,554 | 10,057 |
| Protein residues | 1,310 | 1,209 |
| Ligands | - | |
| B factors (Å2) | | |
| Protein | -190.11 | -144.67 |
| Ligand | | |
| R.M.S. deviations | | |
| Bond lengths (Å) | 0.003 | 0.003 |
| Bond angles (°) | 0.578 | 0.585 |
| Validation | | |
| MolProbity score | 1.92 | 1.75 |
| Clashscore | 9.30 | 9.83 |
| Rotamer outliers (%) | 0 | 0 |
| Ramachandran plot statistics | | |
| Favored (%) | 94.78 | 96.48 |
| Allowed (%) | 4.83 | 3.52 |
| Disallowed (%) | 0 | 0 |

**Video 1. Animation showing the CSA binding mechanism of VAR2CSA**
